# A practical framework for measuring protein oligomerization equilibria by fluorescence correlation spectroscopy

**DOI:** 10.64898/2026.07.08.737283

**Authors:** Dhanashri Rathod, Kaitlyn Parrott, Marcia Levitus

**Author notes:** These authors contributed equally to this work (Dhanashri Rathod, Kaitlyn Parrott).

## Abstract

Protein oligomerization equilibria are central to many biological processes and are often highly sensitive to environmental conditions such as ionic strength, pH, and ligand binding. Quantitative characterization of these equilibria remains experimentally challenging because stable protein complexes frequently dissociate only at concentrations that are difficult to access with conventional biophysical methods. Fluorescence correlation spectroscopy (FCS) is uniquely suited to this problem, as it provides direct access to diffusion coefficients of fluorescently labeled proteins at nanomolar concentrations. However, the quantitative interpretation of FCS data from oligomeric systems requires a rigorous mathematical framework and careful experimental practice that have not previously been described in sufficient detail to guide implementation. Here, we provide a comprehensive description of the experimental workflow and analytical framework for determining dissociation equilibrium constants by FCS, covering instrument calibration, sample preparation, data quality control, after-pulse correction, and nonlinear least-squares fitting. We discuss common sources of error and provide practical guidance on critical experimental considerations including surface passivation, buffer preparation, equilibration time, and the role of labeling efficiency. Using the homotrimeric sliding clamp PCNA as a model system, we demonstrate the complete workflow under a range of KCl concentrations and show that moderate ionic strength stabilizes the PCNA trimer while very high salt partially destabilizes the complex. The approach is general and applicable to any reversible protein self-association reaction accessible by fluorescence detection at low protein concentrations.

## 1 Introduction

Protein self-association into oligomeric complexes is a widespread phenomenon in biology. In this context, oligomerization refers to the reversible association of protein subunits into discrete assemblies with well-defined stoichiometry and quaternary structure, distinguishing it from nonspecific aggregation and other forms of heterogeneous self-association. Approximately two-thirds of human enzymes with known stoichiometry are oligomeric [1], and homo- and hetero-oligomers play essential roles in signal transduction, DNA replication, metabolism, and structural organization of the cell [2]. The equilibrium between oligomeric states often plays a direct functional role, and changes in quaternary structure can profoundly affect protein stability, activity, protein-protein interactions, and downstream signaling pathways. Protein oligomerization is frequently modulated by environmental factors such as ionic strength, pH, phosphorylation, and ligand binding [3–9]. The quantitavie characterization of oligomerization equilibria is therefore an important goal in biophysics and biochemistry.

Characterizing these equilibria requires the quantitative determination of the dissociation equilibrium constant (*K*_*d*_), which sets the concentration scale over which the transition between oligomeric states occurs. The experimental determination of *K*_*d*_ requires measuring an observable sensitive to the degree of dissociation as a function of protein concentration.

This creates a fundamental practical challenge. For stable complexes with *K*_*d*_ values in the micromolar range or below, measurements must be carried out at similarly low protein concentrations to observe significant dissociation. Moreover, because the transition between the fully associated and dissociated states spans several orders of magnitude in concentration, experiments must be designed to cover this entire range reliably. Both requirements place measurements well below the detection limits of most standard biophysical techniques, including analytical ultracentrifugation, isothermal titration calorimetry, and dynamic light scattering [10–12].

Single-molecule fluorescence techniques are particularly well suited to this challenge because of their extraordinary sensitivity, which allows the detection of fluorescently labeled proteins at low nanomolar concentrations. Among these, fluorescence correlation spectroscopy (FCS) has emerged as a powerful and increasingly accessible approach for characterizing protein oligomerization in solution. FCS is based on the statistical analysis of fluorescence intensity fluctuations that arise as labeled molecules diffuse through a femtoliter-sized confocal observation volume. The temporal behavior of these fluctuations, quantified by the autocorrelation function, encodes information about the translational diffusion coefficients of the fluorescent species present in the sample. Because the diffusion coefficient of a globular protein scales approximately with the cube root of its molecular mass, FCS can in principle distinguish between different oligomeric states and report on the degree of dissociation of a complex as a function of concentration, time, or solution conditions.

The application of FCS to protein oligomerization equilibria and kinetics has grown steadily over the past two decades, with successful studies of dimers, trimers, tetramers, and higher-order assemblies [13–18]. Despite this growing body of work, the quantitative analysis of FCS data from oligomeric systems is not straightforward. A key complication is that the autocorrelation decay of a mixture of oligomeric species is experimentally indistinguishable from that of a single species, a consequence of the modest relative differences in diffusion coefficient between, for example, a monomer and its dimer. The measured quantity is therefore an apparent diffusion time (*τ*_app_) that reflects the weighted average of all species present in the sample, where the weighting depends on both the concentrations and the relative brightnesses of each species. Because larger oligomers carry more fluorescent labels and therefore contribute more strongly to the autocorrelation signal, simple intuitive interpretations of *τ*_app_ values can be seriously misleading without a proper mathematical framework.

We previously addressed this problem by deriving a rigorous mathematical treatment that establishes analytical relationships between the experimentally measured *τ*_app_ values and the thermodynamic and kinetic parameters that govern protein oligomerization, including dissociation equilibrium constants, dissociation rate constants, and association rate constants [19]. That work demonstrated that empirical approaches commonly used in the field, such as fitting concentration-dependent *τ*_app_ data with sigmoidal curves or fitting kinetic data with exponential functions, can yield seemingly reasonable results that are in fact inconsistent with the underlying physical model. Physically accurate models are therefore essential for the reliable determination of oligomerization parameters. The mathematical framework developed in that work has since been applied in our laboratory to investigate the oligomerization properties of several biologically important protein systems, including DNA replication sliding clamps [9, 13] and RuBisCO activase [8, 14, 20], where it has yielded quantitative insight into the dependence of complex stability on solution conditions such as ionic strength and osmolyte concentration.

Despite the availability of this mathematical framework, a practical and detailed description of how to implement and analyze FCS experiments for the determination of protein oligomerization equilibria has not been published. The gap between the theoretical treatment and actual experimental practice encompasses a range of important considerations that are rarely discussed in sufficient detail in primary research articles: the design of concentration series relative to the expected *K*_*d*_, the calibration of the confocal observation volume, the acquisition and quality control of autocorrelation data, and the fitting procedures required to extract *K*_*d*_ from the concentration-dependent *τ*_app_ data. Without guidance on these practical aspects, implementing the method from first principles is a significant undertaking that may discourage its adoption by groups that could benefit from it.

Here, we provide a comprehensive description of FCS experiments and data analysis for the determination of protein dissociation equilibrium constants, using the proliferating cell nuclear antigen (PCNA) sliding clamp as a model system. PCNA is a homotrimeric ring-shaped protein that plays a central role in DNA replication and repair by encircling double-stranded DNA and tethering polymerases to their templates [21]. Its oligomeric stability and the dependence of that stability on solution conditions make it an ideal and well-characterized model for demonstrating the method. We describe the complete experimental workflow from sample preparation through data collection and analysis, illustrate each step with data collected at varying KCl concentrations to probe the effect of salt concentration on PCNA stability, and discuss common sources of error and how to identify and avoid them. We anticipate that this practical guide will make quantitative FCS-based oligomerization studies more accessible to the broader biophysics and biochemistry community.

## 2 Fluorescence Correlation Spectroscopy

### 2.1 Principles and Instrumentation

Fluorescence correlation spectroscopy (FCS) is a single-molecule sensitive technique that extracts information about molecular dynamics from the statistical analysis of fluorescence intensity fluctuations. In a typical FCS experiment, a laser beam is focused through a high-numerical-aperture objective to create a diffraction-limited confocal observation volume of femtoliter dimensions (Figure 1). Fluorescent molecules diffusing through this volume give rise to bursts of emitted photons, and the resulting fluctuations in detected fluorescence intensity carry information about the concentration and diffusion coefficients of the fluorescent species in solution. These fluctuations are small when the average number of fluorescent molecules in the observation volume is large, and become more pronounced as the concentration decreases toward the single-molecule regime. For this reason, FCS experiments are typically performed at concentrations in the low nanomolar range, which as we discuss below is precisely the concentration range required to observe the dissociation of stable protein complexes.

**Figure 1:**
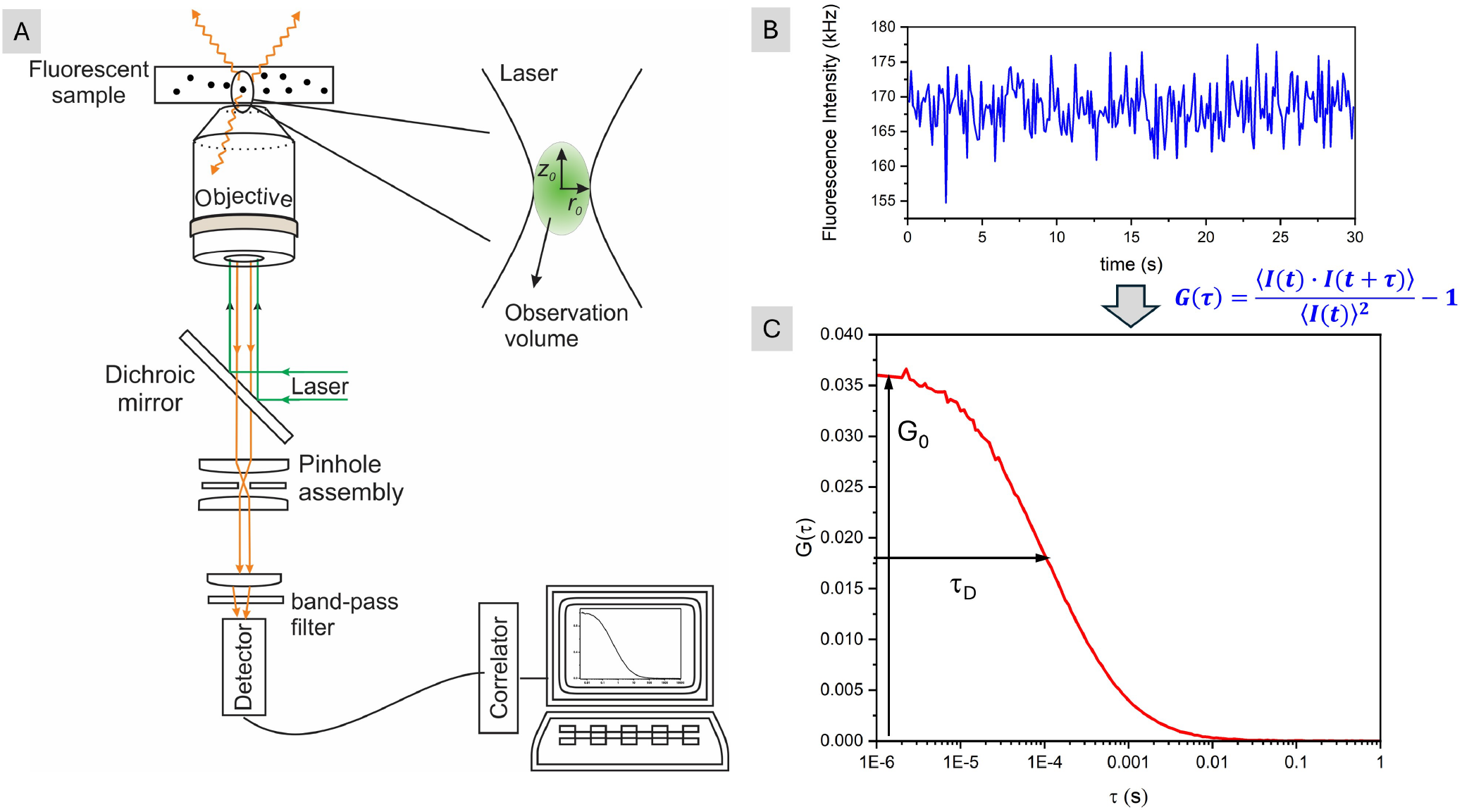
Schematic of a confocal fluorescence correlation spectroscopy (FCS) instrument. (A) The excitation laser beam is directed through a dichroic mirror into a high-numerical-aperture oil-immersion objective, which focuses it to a diffraction-limited confocal observation volume of femtoliter dimensions, characterized by radial and axial semiaxes *r*_0_ and *z*_0_, respectively. Fluorescence emitted by labeled molecules diffusing through this volume is collected through the same objective, separated from the excitation light by the dichroic mirror, spatially filtered by a pinhole, spectrally filtered by a bandpass filter, and detected by an avalanche photodiode. The resulting photon counts are passed to a hardware correlator, which computes the autocorrelation function in real time. (B) Representative fluorescence intensity trace *I*(*t*) recorded from a 10 nM solution of TMR in water. (C) Autocorrelation function *G*(*τ*) computed from the intensity trace according to Equation 1. The amplitude *G*_0_ is inversely proportional to the average number of fluorescent particles in the observation volume, and the decay time *τ*_*D*_ reports on the translational diffusion coefficient of the fluorescent species.

The quantity of interest in an FCS experiment is the time-dependent fluorescence intensity *I*(*t*), which fluctuates around its mean value ⟨*I*⟩ as molecules diffuse into and out of the observation volume. The temporal behavior of the fluorescence fluctuations is quantified by the autocorrelation function *G*(*τ*), defined as

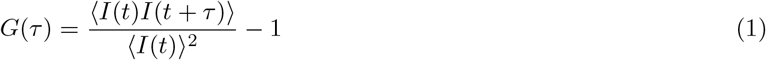

where *τ* is the lag time and angular brackets denote time averages over the duration of the measurement. A representative fluorescence intensity trace recorded from a 10 nM solution of TMR in water and the corresponding autocorrelation decay are shown in Figure 1. Conceptually, *G*(*τ*) measures the self-similarity of the fluorescence signal at two time points separated by a lag time *τ*. The fluorescence intensity detected from a single molecule depends on its position within the observation volume, being highest when the molecule is near the focal point where both excitation and detection efficiency are greatest. As molecules diffuse, their positions change and the detected intensity fluctuates accordingly. At short lag times, where the root-mean-square displacement is small, a molecule remains close to its position at time *t*, so *I*(*t*) and *I*(*t* + *τ*) are similar and *G*(*τ*) is large. As *τ* increases and molecules diffuse further from their initial positions, the intensity at time *t* + *τ* becomes progressively less correlated with that at time *t*, and *G*(*τ*) decays to zero. The characteristic time over which this decay occurs, the diffusion time *τ*_*D*_, is therefore a measure of how long a fluorescent particle takes, on average, to traverse the observation volume, which in turn depends on its translational diffusion coefficient *D*.

In principle, *G*(*τ*) could be computed from a recorded *I*(*t*) trace in post-processing. This approach requires time-correlated single photon counting (TCSPC) hardware and a pulsed laser to achieve the nanosecond time resolution needed to capture the earliest lag times of the autocorrelation decay, and is the basis of techniques such as fluorescence lifetime correlation spectroscopy (FLCS) [22]. A simpler and less expensive alternative, which is the most widely implemented form of FCS and the approach used in this work, is to use a continuous-wave laser and a hardware correlator card that computes *G*(*τ*) in real time as photons are detected. The output of the experiment is therefore the autocorrelation function itself rather than the raw intensity signal, and it is *G*(*τ*) that is analyzed to extract the diffusion time *τ*_*D*_ and, from it, information about the size and concentration of the fluorescent species in solution.

Modern FCS implementations can extract considerably more information than the autocorrelation function alone. Techniques such as fluorescence lifetime correlation spectroscopy (FLCS) [22], two-color fluorescence cross-correlation spectroscopy (FCCS) [23], FCS-FRET [24], and photon counting histogram analysis (PCH) [25] provide additional observables that can complement or extend the analysis described here. However, these approaches require more sophisticated instrumentation and data analysis, and are beyond the scope of this work. Throughout this paper, we assume that the only measured quantity is the autocorrelation function *G*(*τ*), computed using a continuous-wave laser and a hardware correlator as described above, which is sufficient for the quantitative determination of *K*_*d*_ as described in the following sections.

For a monodisperse solution of freely diffusing fluorescent particles, the autocorrelation function takes the form [26, 27]

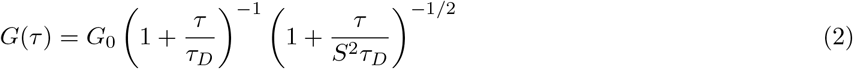

where *G*_0_ is the amplitude of the autocorrelation function at *τ* = 0, 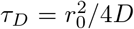 is the characteristic diffusion time, *r*_0_ is the radial semiaxis of the Gaussian observation volume, and *S* = *z*_0_*/r*_0_ is the geometrical parameter defined as the ratio of the axial semiaxis *z*_0_ to the radial semiaxis *r*_0_ (Figure 1). The amplitude *G*_0_ is inversely proportional to the average number of fluorescent particles in the observation volume, ⟨*N*⟩, and therefore provides an independent measure of concentration

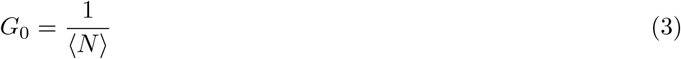

where ⟨*N*⟩ = *CN*_*A*_*V*_eff_ is determined by the molar concentration of fluorescent particles *C*, Avogadro’s number *N*_*A*_, and the effective observation volume 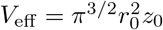 [26, 27].

In a typical FCS instrument, *S* is in the range of 5 to 10, meaning that the observation volume is significantly elongated along the optical axis. When *S* ≫ 1, the condition *τ/*(*S*^2^*τ*_*D*_) ≪ 1 holds over the entire range of lag times where the first term in Equation 2 varies appreciably, and the second term contributes negligibly to the decay. Equation 2 then simplifies to

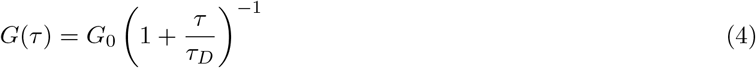

Physically, this simplification reflects the fact that when the observation volume is highly elongated, axial diffusion contributes negligibly to fluorescence fluctuations. The autocorrelation decay is therefore governed almost entirely by the lateral component of diffusion, and Equation 4 is mathematically equivalent to the autocorrelation function expected for two-dimensional diffusion in the radial plane. Its practical importance is that it allows the derivation of fully analytical relationships between the experimentally measured *τ*_app_ values and the dissociation equilibrium constant *K*_*d*_, as described in Section 3.

### 2.2 Diffusion Coefficients of Oligomeric Proteins

The translational diffusion coefficient *D* of a globular protein in solution is related to its hydrodynamic radius *R*_*h*_ through the Stokes-Einstein equation,

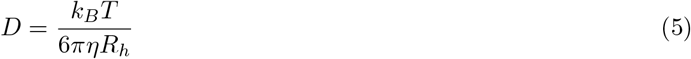

where *k*_*B*_ is Boltzmann’s constant, *T* is the absolute temperature, and *η* is the solvent viscosity. For a globular protein, the hydrodynamic radius *R*_*h*_ scales approximately as the cube root of the molecular volume. Since molecular volume is approximately proportional to molecular mass *M*, *R*_*h*_ ∝ *M* ^1*/*3^. Thus, for an oligomer composed of *n* identical subunits, *R*_*h*_ ∝ *n*^1*/*3^. As a result, the diffusion coefficient of an *n*-mer is related to the monomer diffusion coefficient *D*_1_ as

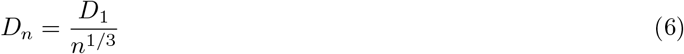

and the corresponding diffusion times scale as

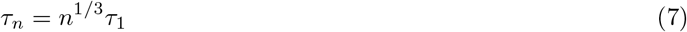

where *τ*_1_ is the diffusion time of the monomer. This weak dependence on oligomeric state has an important practical consequence: the diffusion times of different oligomeric species lie within a relatively narrow range. For the trimeric protein PCNA studied in this work, the intact trimer is expected to diffuse only 3^1*/*3^ ≈ 1.44 times more slowly than its dissociated monomers, corresponding to a difference in diffusion time of 44%. For a dimeric protein the difference is even smaller, with the dimer diffusing only 2^1*/*3^ ≈ 1.26 times more slowly than the monomer. These modest differences mean that FCS cannot resolve individual oligomeric species directly from the shape of the autocorrelation decay, but as we describe in the following section, the protein concentration dependence of the apparent diffusion time *τ*_app_ contains quantitative information about the distribution of oligomeric states in solution and can be used to determine *K*_*d*_ provided an appropriate mathematical framework is applied.

### 2.3 FCS of Oligomeric Proteins

For a solution containing a mixture of fluorescent species, where each species is defined as a population of particles with the same diffusion coefficient and brightness, the total autocorrelation function is given by [19]

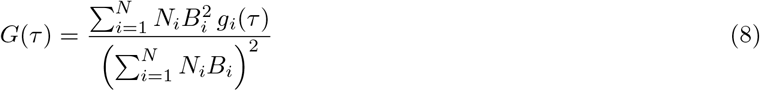

where *N*_*i*_ is the average number of molecules of species *i* in the observation volume, and *B*_*i*_ is the molecular brightness of species *i*. The function *g*_*i*_(*τ*) is the normalized autocorrelation function of a homogeneous solution of species *i*, defined such that its amplitude equals unity (*G*_0_ = 1 in Equation 4): *g*(*τ*) = (1 + *τ/τ*_*D*_)^−1^. The weighting of each term in the numerator by 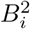 reflects the fact that brighter particles produce larger intensity fluctuations and therefore contribute more strongly to the autocorrelation signal.

To determine the dissociation equilibrium constant *K*_*d*_ under a given set of solution conditions, FCS measurements are performed as a function of protein concentration (*C*_0_) to obtain a titration curve of *τ*_app_ vs. *C*_0_. At high protein concentrations, where dissociation is negligible, the solution is dominated by the intact oligomer and *τ*_app_ approaches the characteristic diffusion time of the oligomer, *τ*_*m*_, where *m* is the oligomeric stoichiometry (e.g. *m* = 2 for a dimer). At low protein concentrations, dissociation becomes significant and a mixture of oligomeric species coexists in solution, causing *τ*_app_ to shift toward the diffusion time of the monomer, *τ*_1_. The shape of the resulting titration curve contains quantitative information about *K*_*d*_, as described in Section 3.

A practical complication arises at high protein concentrations, where the number of fluorescent particles in the observation volume becomes too large for reliable FCS measurements. To circumvent this, titration experiments are typically performed using a fixed low concentration of fluorescently labeled protein *C*_*L*_, diluted into variable concentrations of unlabeled protein *C*_*U*_. Since labeling is rarely complete, *f C*_*L*_ represents the actual concentration of labeled subunits in solution, where *f* is the labeling efficiency (0 *< f* ≤ 1). FCS measurements are most reliable when the average number of fluorescent particles in the observation volume, ⟨*N*⟩, is in the range of approximately 1 to 100, which for a typical observation volume of a few femtoliters corresponds to labeled protein concentrations in the nanomolar range. By fixing *C*_*L*_ at a low nanomolar concentration, the amplitude of the autocorrelation function remains within this optimal range throughout the titration, while the total protein concentration *C*_0_ = *C*_*L*_ + *C*_*U*_, which governs the degree of dissociation, can be varied over several orders of magnitude by adding increasing amounts of unlabeled protein.

Throughout this work, *C*_0_, *C*_*L*_, and *C*_*U*_ represent protein concentrations expressed in monomer equivalents, which are the natural variables for the equilibrium and mass-balance relationships developed in the following sections. When referring to concentrations in terms of oligomers, we use the same symbols with a tilde: 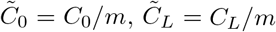, and 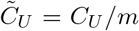, where *m* is the oligomeric stoichiometry (*m* = 3 for PCNA). For ease of interpretation, figures are plotted as a function of PCNA trimer concentration 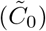 on the bottom axis, with the corresponding monomer concentration, *C*_0_, shown on the top axis. The two are related through 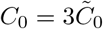.

It is convenient to define *p* = *f C*_*L*_*/C*_0_ as the fraction of total protein subunits in solution that carry a fluorescent label, a quantity that appears throughout the analysis (Figure 2). As a consequence of mixing labeled and unlabeled protein, oligomers of the same stoichiometry *n* may carry different numbers of fluorescent labels *b*, where 0 ≤ *b* ≤ *n*, depending on how many of their subunits were drawn from the labeled stock. These species have the same diffusion coefficient but different brightnesses, and must therefore be treated as distinct species in Equation 8.

**Figure 2:**
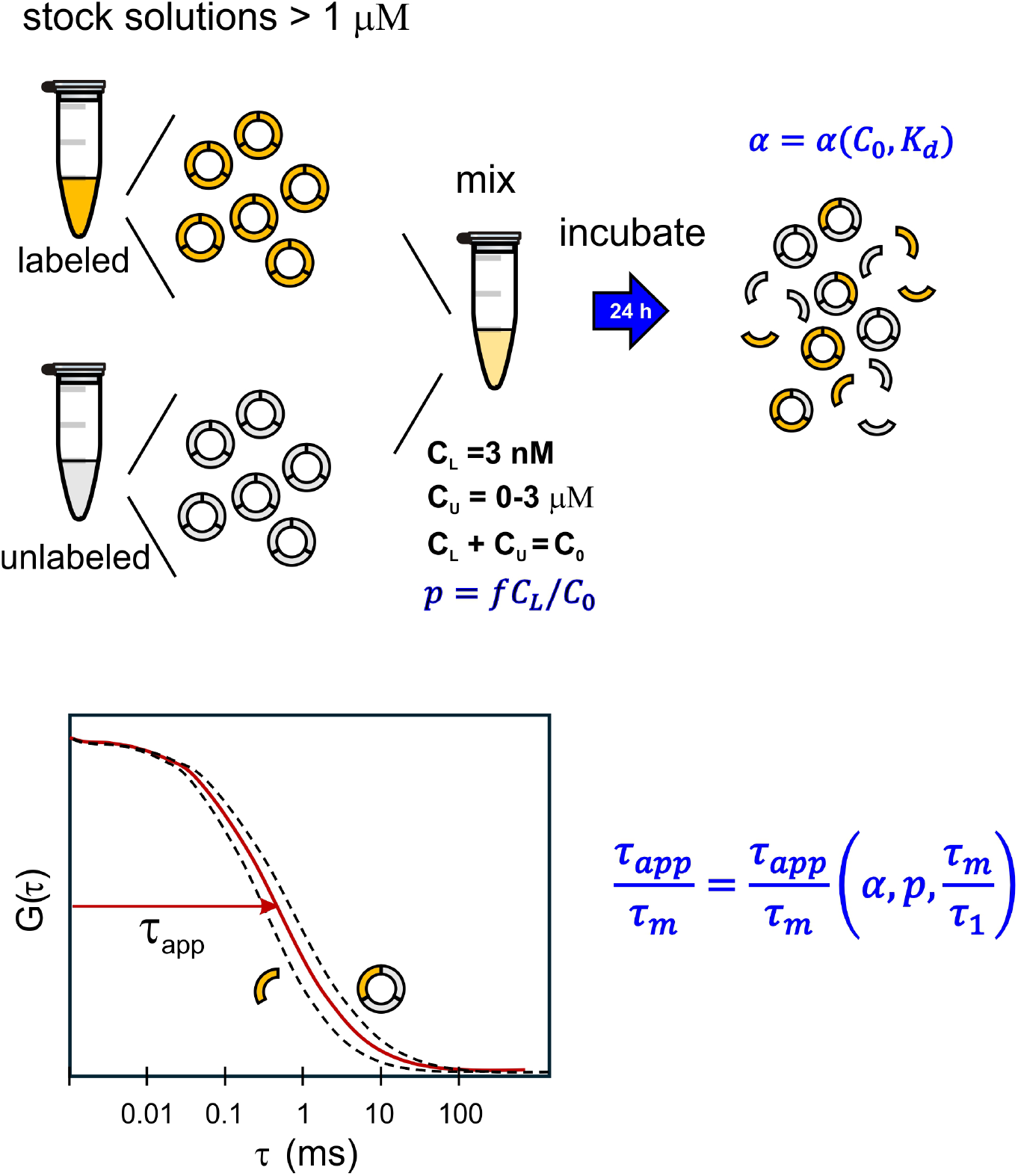
Experimental Strategy: Experiments are performed with a fixed concentration of labeled protein 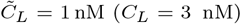 and a variable concentration of unlabeled protein 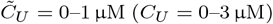. The fraction of labeled subunits *p* = *f C*_*L*_*/C*_0_ decreases as *C*_0_ increases. Oligomers of the same stoichiometry *n* may carry between 0 and *n* fluorescent labels and are treated as distinct species in the analysis. The degree of dissociation, *α*, depends on the total protein concentration, *C*_0_ and the thermodynamic dissociation constant, *K*_*d*_.The apparent diffusion time *τ*_app_ is obtained by fitting the experimental autocorrelation function with Equation 4. Its value depends on *τ*_1_ and *τ*_*m*_, the diffusion times of the monomer and intact oligomer respectively, the degree of dissociation *α*, which sets the relative populations of monomers and oligomers, and the labeling fraction *p*, which determines the distribution of fluorescent labels among oligomers of the same stoichiometry and therefore their relative brightnesses.

Letting *N*_*n,b*_ denote the average number of oligomers with *n* subunits and *b* fluorescent labels present in the observation volume, and assuming that the brightness of each oligomer is directly proportional to the number of labels it carries, Equation 8 can be rewritten as

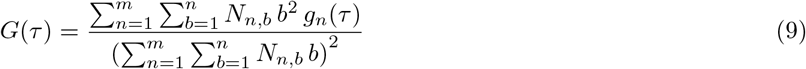

Note that *g*_*n*_(*τ*) depends only on *n* and not on *b*, since oligomers with the same stoichiometry are assumed to diffuse at the same rate regardless of how many labels they carry. The distribution of *b* for a given *n* follows a binomial distribution with parameter *p*, since each subunit in an oligomer is independently drawn from the pool of labeled and unlabeled protein with probability *p* of carrying a fluorescent label. The derivation of the analytical expressions that result from this binomial distribution is provided in the Supporting Information of Kanno and Levitus [19], where it is shown that Equation 9 reduces to

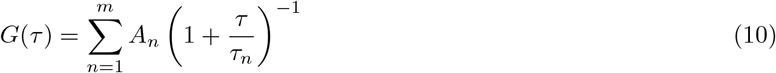

where

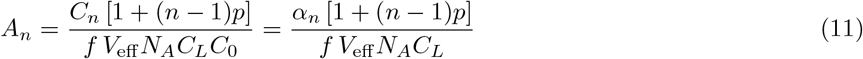

The coefficients *A*_*n*_ represent the contributions of the *n*-mer to the total autocorrelation function and *C*_*n*_ is the molar concentration of each *n*-mer, expressed in terms of monomers. In the second expression, *α*_*n*_ = *C*_*n*_*/C*_0_ is the fractional concentration of the *n*-mer, defined as the fraction of total protein subunits that are part of an *n*-mer, which satisfies 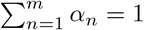.

It is convenient to define the prefactor

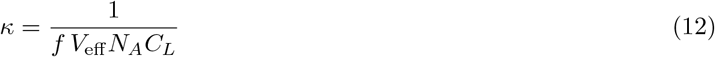

which depends only on instrumental parameters (*V*_eff_), the concentration of labeled protein (*C*_*L*_), and the labeling efficiency (*f*). In our experiments, *C*_0_ is varied by adding increasing amounts of unlabeled protein while keeping *C*_*L*_ fixed, so *κ* remains constant throughout the titration. With this definition, Equation 11 simplifies to

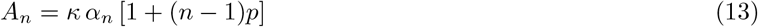

The concentration dependence of *A*_*n*_ enters entirely through *α*_*n*_ = *C*_*n*_*/C*_0_ and *p* = *f C*_*L*_*/C*_0_, both of which vary with *C*_0_: *α*_*n*_ reflects how the equilibrium distribution of oligomeric species shifts with total protein concentration, while *p* decreases as *C*_0_ increases because the labeled subunits become an ever smaller fraction of the total protein in solution.

A key practical consequence of Equation 10 is that the autocorrelation decay of a mixture of oligomers is experimentally indistinguishable from the decay of a single species. This is a direct result of the weak dependence of *τ*_*n*_ on *n*: because diffusion times scale as *n*^1*/*3^, the individual contributions of different oligomeric species to *G*(*τ*) overlap substantially and cannot be resolved from the shape of the decay alone. Experimental autocorrelation decays are therefore fitted with the single-species form of Equation 4 to recover an apparent diffusion time *τ*_app_, which satisfies *τ*_1_ ≤ *τ*_app_ ≤ *τ*_*m*_.

We note that because of the brightness-squared weighting, *τ*_app_ is not a simple linear average of the diffusion times of the individual species weighted by their molar concentrations.

The fractional concentrations *α*_*n*_ are not independent variables but are determined by *K*_*d*_ and *C*_0_ through the equilibrium expressions and mass balance conditions described in Section 3. For a given value of *K*_*d*_, the concentrations of all oligomeric species at each *C*_0_ are fully determined, and therefore so are the amplitudes *A*_*n*_ and the predicted *τ*_app_. The experimental titration curve of *τ*_app_ vs. *C*_0_ can therefore be fitted with *K*_*d*_ as the only free parameter, provided that *C*_*L*_, *f*, and *τ*_*m*_ are known independently. Here *τ*_*m*_ is the diffusion time of the intact oligomer, determined experimentally in conditions where dissociation is negligible and *τ*_app_ ≈ *τ*_*m*_ (Section 4.5). The fitting procedure is described in detail in Section 3.The key quantities governing *τ*_app_ and their relationships are summarized in Figure 2.

## 3 Analysis

The mathematical framework developed by Kanno and Levitus [19] was derived for the general case of oligomeric proteins but was illustrated specifically for dimeric and tetrameric proteins. Here, we adapt the framework for a homotrimeric protein, as required for the analysis of FCS experiments performed with PCNA. Previous studies have provided evidence that PCNA dissociates cooperatively, with no detectable accumulation of dimeric intermediates, so that the equilibrium can be described as a two-state transition between the intact trimer and free monomers [13].

### 3.1 Equilibrium Expression

The degree of dissociation *α* at a given total protein concentration 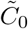 is determined by the dissociation equilibrium constant *K*_*d*_ through the equilibrium expression and mass balance for the specific oligomerization reaction under consideration. In the equations below, *C*_0_ is expressed in terms of monomers, i.e. 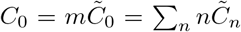, so that the mass balance accounts for the fact that each oligomer of stoichiometry *n* contains *n* subunits. The resulting relationship *α* = *α*(*K*_*d*_, *C*_0_) is analytical for the most common oligomerization mechanisms, as shown below.

For a homodimeric protein dissociating into monomers, D ⇌ 2M, the dissociation equilibrium constant is *K*_*d*_ = [M]^2^*/*[D], and the mass balance gives *C*_0_ = [M] + 2[D]. The degree of dissociation *α* = [M]*/C*_0_ is the positive root of the quadratic equation

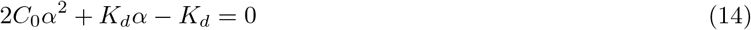

which gives *α* as a function of two variables: *C*_0_ and *K*_*d*_. The analytical expression for *α*(*C*_0_, *K*_*d*_) is given in the Supplemental Information file (Equation S1).

For a homotrimeric protein dissociating into monomers, T ⇌ 3M, which is the case relevant to PCNA studied in this work, the dissociation equilibrium constant is *K*_*d*_ = [M]^3^*/*[T] and the mass balance gives *C*_0_ = [M] + 3[T], where *C*_0_ is expressed in terms of monomers as before. Defining the degree of dissociation as *α* = [M]*/C*_0_, so that [M] = *αC*_0_ and [T] = (1 − *α*)*C*_0_*/*3, and substituting into the expression for *K*_*d*_ gives

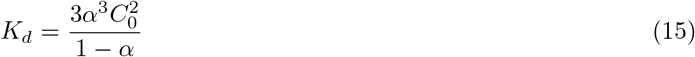

or equivalently,

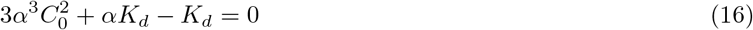

Note that *α* corresponds to *α*_1_ = *C*_1_*/C*_0_ as defined in Section 2, and 1−*α* corresponds to *α*_3_ = *C*_3_*/C*_0_ = 3[T]*/C*_0_, so that *α*_1_ + *α*_3_ = 1 as required. Equation 16 has only one positive real root (Equation S2), which gives the degree of dissociation as a function of the total protein concentration and the dissociation equilibrium constant (*α* = *α*(*C*_0_, *K*_*d*_), see Supplemental Information file).

In general, regardless of the oligomerization mechanism, the degree of dissociation, *α* = *α*(*C*_0_, *K*_*d*_), is a function of only two variables: the total protein concentration and the dissociation equilibrium constant. This function plays a central role in the analysis: for a given value of *K*_*d*_, it determines the distribution of oligomeric species at each concentration point in the titration, and therefore the predicted value of *τ*_app_ at that concentration, as described in the following section.

### 3.2 Relating *τ*_app_ to *K*_*d*_

For a mixture containing only monomers and a single oligomeric species of stoichiometry *m*, whether a dimer (*m* = 2) or a trimer (*m* = 3), Equation 10 reduces to two terms,

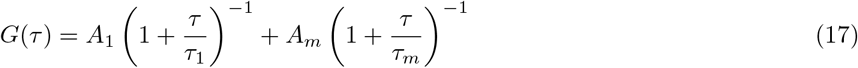

where *τ*_*m*_ = *m*^1*/*3^*τ*_1_ is the diffusion time of the oligomer and *A*_1_ and *A*_*m*_ are the amplitudes defined in Equation 13.

As established in Section 2, the autocorrelation decay of a mixture of monomers and trimers is experimentally indistinguishable from that of a single species, and it is therefore not possible to resolve *τ*_1_ and *τ*_*m*_ individually from the shape of the decay. Experimental autocorrelation decays are instead fitted with the single-species form of Equation 4 to recover a single quantity, *τ*_app_, whose concentration dependence encodes information about the equilibrium between monomers and oligomers.

Kanno and Levitus [19] derived the analytical expression that relates *τ*_app_ to *τ*_1_, *τ*_*m*_, and the degree of dissociation *α*. This dependence on *α* is physically intuitive: when *α* = 1 the solution contains only monomers and *τ*_app_ = *τ*_1_, and when *α* = 0 it contains only trimers and *τ*_app_ = *τ*_*m*_. At intermediate values of *α, τ*_app_ satisfies *τ*_1_ ≤ *τ*_app_ ≤ *τ*_*m*_ and is given as the positive root of the quadratic equation

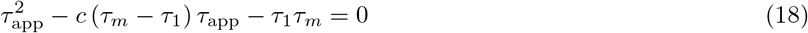

where

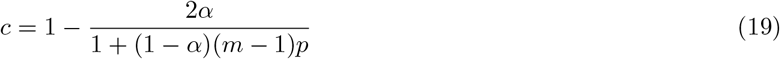

and *p* = *f C*_*L*_*/C*_0_ is the labeling fraction defined in Section 2. The original derivation in Kanno and Levitus was for the specific case of a homodimer (*m* = 2), where Equation 19 reduces to *c* = 1 − 2*α/*(1 + (1 − *α*)*p*). The derivation of the generalized expression for arbitrary oligomeric stoichiometry *m*, which follows the same approach as in Kanno and Levitus [19], is provided in the Supplemental Information file.

The positive root of Equation 18 is

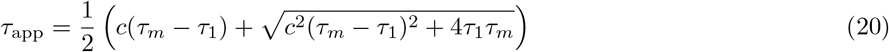

Together with Equation S1 for a dimer or Equation S2 for a trimer, Equation 20 establishes the quantitative relationship between the experimentally measured *τ*_app_ and the dissociation equilibrium constant *K*_*d*_. The fitting procedure that exploits this relationship to extract *K*_*d*_ from the experimental titration curve is described in Section 3.4.

### 3.3 Fitting of Normalized *τ*_app_ Values

In principle, the fitting requires independent knowledge of both *τ*_1_ and *τ*_*m*_ as inputs. However, it is convenient in practice to work with normalized apparent diffusion times, defined as *τ*_app_*/τ*_*m*_, where *τ*_*m*_ is the diffusion time of the intact oligomer, measured experimentally in conditions where dissociation is negligible so that *τ*_app_ ≈ *τ*_*m*_ (section 4.5). In terms of this normalized quantity, Equation 20 becomes:

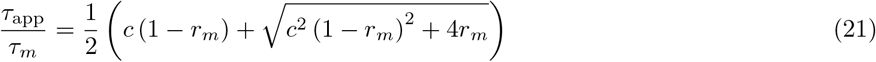

which depends on *τ*_1_ and *τ*_*m*_ only through their ratio *r*_*m*_ = *τ*_1_*/τ*_*m*_. This formulation leads naturally to two fitting strategies. In the first, *r*_*m*_ is treated as a free parameter alongside *K*_*d*_, resulting in a two-parameter fit. In the second strategy, *r*_*m*_ is fixed at the value *m*^−1*/*3^ predicted by Equation 7 for a globular protein, reducing the fit to a single free parameter *K*_*d*_. The validity of this approach rests on the assumption that the hydrodynamic radii of the monomer and trimer scale as expected for a globular particle. In either case, *τ*_*m*_ is determined experimentally by measuring *τ*_app_ immediately after dilution from a concentrated stock, before significant dissociation has occurred. For stable oligomers such as PCNA, the slow dissociation kinetics ensure that *τ*_app_ ≈ *τ*_*m*_ on the timescale of the measurement, providing a reliable estimate of *τ*_*m*_. The normalized titration curve *τ*_app_*/τ*_*m*_ vs. PCNA concentration is then fitted to extract *K*_*d*_. The experimental determination of *τ*_*m*_ is illustrated in Figure 3.

**Figure 3:**
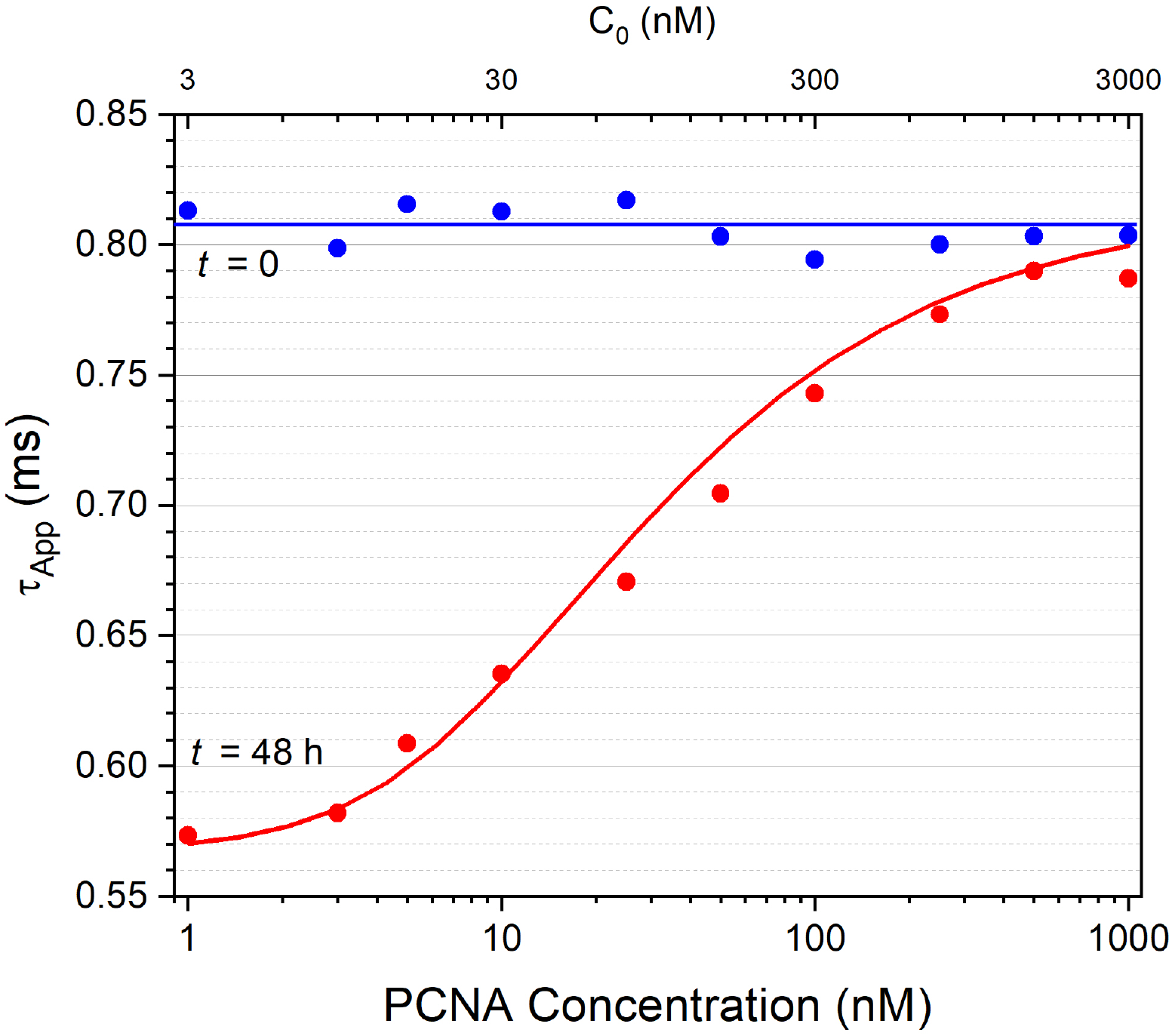
Experimental determination of *τ*_3_, the diffusion time of the intact PCNA trimer. Blue circles show *τ*_app_ measured immediately after dilution from a concentrated stock, before significant dissociation has occurred, so that *τ*_app_ ≈ *τ*_3_ at all concentrations. The horizontal blue line shows the mean value used as *τ*_3_ in the normalization of the titration curves. Red circles show *τ*_app_ measured after 48 hours of equilibration at each PCNA concentration, and the solid red curve shows the best fit to the equilibrated data. Note that unlike the titration curves shown in Section 5, the y-axis here shows absolute values of *τ*_app_ in ms rather than the normalized ratio *τ*_app_*/τ*_3_. The bottom axis shows PCNA concentration expressed in terms of trimers 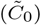; the top axis shows the corresponding total monomer concentration *C*_0_. The systematic decrease in *τ*_app_ at low PCNA concentrations reflects dissociation of the trimer into monomers at equilibrium.

**Figure 4:**
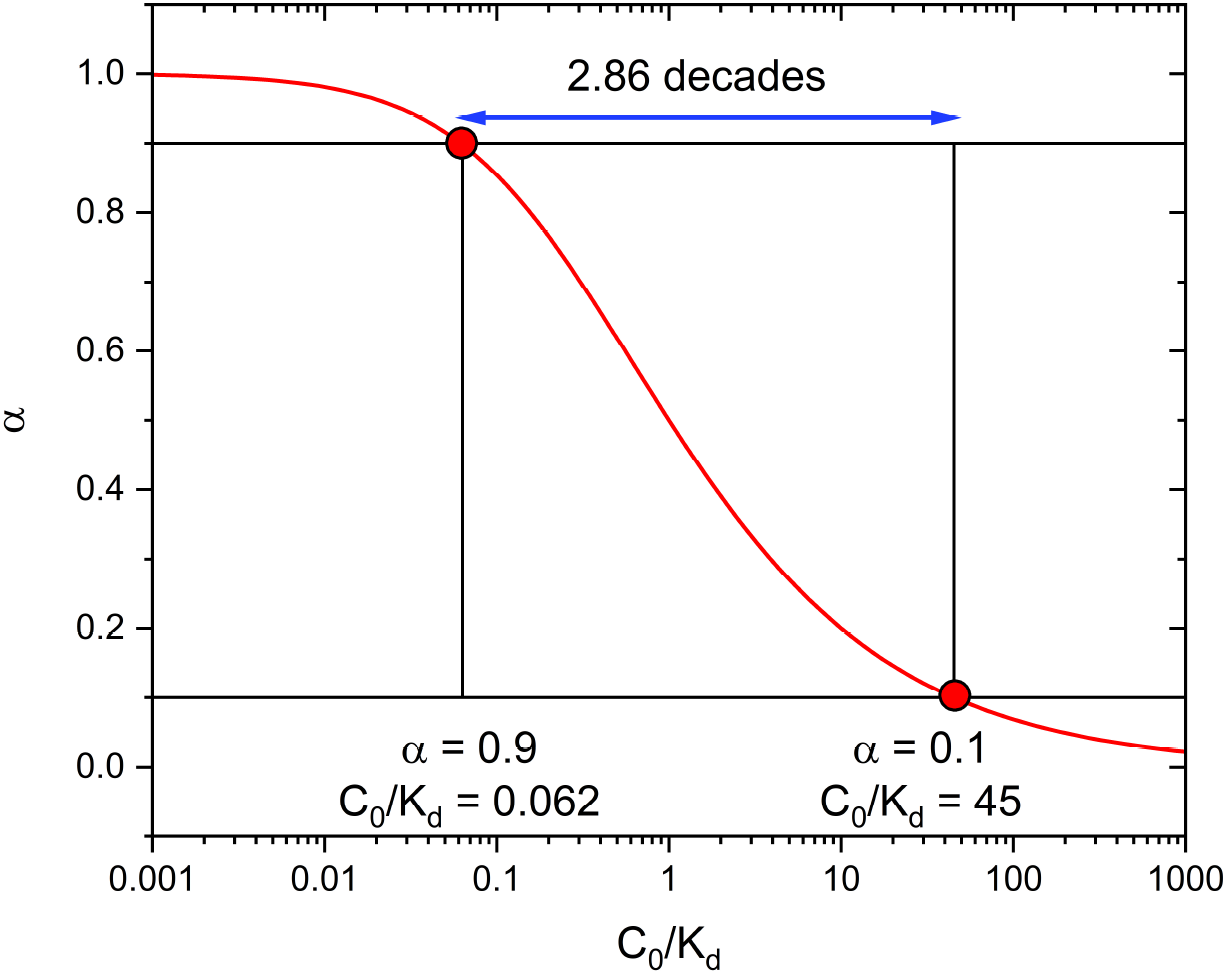
Logarithmic span of a homodimer dissociation equilibrium. The fractional concentration of dimer (*α*) is shown as a function of the reduced protein concentration *C*_0_*/K*_*d*_. The logarithmic span, defined as the range of *C*_0_*/K*_*d*_ over which *α* changes from 0.9 to 0.1, spans 2.86 decades for a homodimer (blue arrow). The corresponding values of *C*_0_*/K*_*d*_ are 0.062 and 45, respectively.

This normalization strategy was employed in the experiments described in this work and has an important practical advantage. The diffusion time 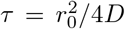 is not a molecular property of the protein but depends on the size of the confocal observation volume through *r*_0_, which can vary from day to day due to small changes in laser alignment. As a result, absolute values of *τ*_app_ measured on different days or on different instruments are not directly comparable. The normalized ratio *τ*_app_*/τ*_*m*_, however, is insensitive to these instrumental factors because both *τ*_app_ and *τ*_*m*_ are measured under identical conditions and depend on *r*_0_ in the same way, so that *r*_0_ cancels in the ratio. The normalized titration curve therefore reflects only the molecular properties of the protein, specifically the equilibrium between monomers and trimers, and can be compared directly across experiments performed on different days or in different laboratories.

### 3.4 Fitting Algorithm

The fitting procedure is implemented as follows. For a given trial value of *K*_*d*_, the degree of dissociation *α* at each measured total protein concentration *C*_0_ is obtained from Equation S1 for a dimer, or Equation S2 for a trimer. The labeling fraction *p* = *f C*_*L*_*/C*_0_ is then evaluated at the same concentration point. With *α* and *p* determined, the coefficient *c* is calculated from Equation 19, and the predicted normalized apparent diffusion time *τ*_app_*/τ*_*m*_ is obtained from Equation 21. This procedure is repeated at each value of *C*_0_ in the titration series to generate the complete predicted curve.

The trial value of *K*_*d*_ is varied and the entire procedure repeated until the predicted curve provides the best fit to the experimental *τ*_app_*/τ*_*m*_ vs. protein concentration data in a least-squares sense. The minimization is performed using standard nonlinear least-squares algorithms, such as the Levenberg-Marquardt method as implemented in software packages including MATLAB or Python. The required experimental inputs are the titration data *τ*_app_*/τ*_*m*_ vs. *C*_0_, the labeled protein concentration *C*_*L*_, and the labeling efficiency *f*. As discussed in Section 5, the extracted *K*_*d*_ is relatively insensitive to the precise value of *f*, so an approximate determination of *f* is sufficient for reliable results.

## 4 Materials and Methods

### 4.1 Instrumentation

The data presented here were acquired using a home-built confocal FCS setup. The output of a 532-nm continuous-wave laser (Compass 215M-10, Coherent) was expanded, collimated, and directed through a dichroic filter into an Olympus PlanApo 100 × /1.4 NA oil-immersion objective. Samples were placed into 50 µL perfusion chambers pretreated with bovine serum albumin (BSA) to minimize protein adsorption onto the cover glass. Fluorescence was collected through the same objective, separated from the excitation light through the dichroic filter, and focused through a 50 µm pinhole onto an avalanche photodiode detector (SPCM-AQR14, Perkin-Elmer Optoelectronics). A bandpass filter (3RD560-620, Omega) was placed before the detector to minimize background signal. Autocorrelation functions were computed in real time using a hardware correlator (ALV5000/EPP USB-25, ALV GmbH, Germany).

Commercial FCS systems are also available from several manufacturers and are equally suitable for the experiments described here.

### 4.2 Protein Preparation and Labeling

The mathematical framework described in sections 2 and 3 assumes that fluorescent labels are introduced at a single defined site per subunit, so that each subunit carries at most one fluorescent label. The labeling efficiency *f*, defined as the fraction of subunits in the labeled stock that carry a fluorescent label, quantifies how completely this site is occupied and is explicitly accounted for in the analysis through the parameter *p* = *f C*_*L*_*/C*_0_ introduced in Section 2. In practice, site-specific labeling with a single label per subunit is achieved by introducing a single reactive group, typically a cysteine residue, at a defined position on the protein surface and reacting it with a maleimide derivative of the desired fluorophore.

The choice of fluorophore is an important practical consideration. The fluorophore must be photostable enough to withstand the laser illumination during data acquisition, and its photophysical properties must be compatible with the assumptions underlying the autocorrelation analysis. In particular, processes that cause transient interruptions in fluorescence emission, such as triplet-state blinking and photoisomerization, give rise to additional fast components in the autocorrelation decay that can interfere with the diffusion-based analysis if not properly accounted for [28–30]. In the experiments described in this work, we use tetramethylrhodamine (TMR), a rhodamine dye that is well characterized photophysically [28, 29, 31]. Under our experimental conditions, no contributions from triplet blinking or other photophysical processes are observed in the autocorrelation decays [29], and no corrections beyond the single-species diffusion model of Equation 4 are required. Other fluorophores commonly used in FCS experiments, including Alexa Fluor dyes and ATTO dyes, are equally suitable provided their photophysical behavior is characterized under the specific experimental conditions used [28].

The labeling efficiency is typically determined from absorbance measurements by calculating the ratio of fluorophore concentration to protein subunit concentration, using the known molar absorptivities of the fluorophore and the protein at their respective absorption maxima and correcting for the contribution of the fluorophore to the absorbance at the protein absorption maximum [32]. As discussed in Section 5, the extracted *K*_*d*_ is relatively insensitive to the precise value of *f*.

As with any fluorescence-based experiment, control experiments should be performed to verify that the fluorescent label does not perturb the properties of the protein under study. Although negative results are rarely reported in the literature, there are documented cases where fluorescent labels affect protein conformation, protein-protein interactions, and other structural and dynamic properties [33–36]. Researchers should therefore characterize labeled proteins as thoroughly as possible to confirm that they behave as their unlabeled counterparts. Depending on the protein system, appropriate controls may include enzymatic activity assays, binding assays with known interaction partners, circular dichroism spectroscopy, melting temperatures, and NMR or X-ray structures whenever possible. For the PCNA experiments described in this work, control experiments confirming that the mutations and labeling do not affect the oligomeric and functional properties of PCNA are reported in [13].

Protein concentrations were determined using a Bradford assay with unlabeled wild-type PCNA as a standard, as described previously [13]. The labeling efficiency *f* was determined by measuring the TMR concentration of the labeled protein by absorbance spectroscopy at 550 nm using a molar absorptivity of *ε* = 91,000 M^−1^cm^−1^, and comparing the TMR concentration to the monomer concentration,

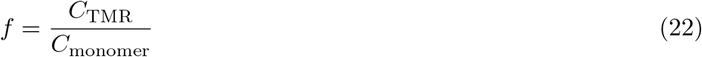

For the PCNA samples used in this work, the labeling efficiency was determined to be *f* = 0.6 ± 0.1.

### 4.3 Chamber Preparation

Non-specific adsorption of protein onto the cover glass surface is a common problem in FCS experiments, as it reduces the concentration of freely diffusing protein in solution and can introduce artifacts in the autocorrelation decay. The appropriate surface passivation strategy depends on the protein under study, and a variety of protocols have been described in the literature, including coating with BSA, polyethylene glycol (PEG), lipid bilayers, or casein [37,38]. Researchers should verify that the chosen passivation strategy is effective for their specific protein by confirming that the count rate and *G*_0_ remain stable over the duration of the experiment. A progressive decrease in count rate accompanied by an increase in *G*_0_ is a clear indication that protein is being lost to adsorption onto the surface, reducing the concentration of freely diffusing labeled molecules in solution. If this is observed, the passivation protocol should be revised before proceeding with the titration experiments.

For the PCNA experiments described in this work, glass slides were cleaned by treatment in an ozonator chamber for 20 minutes (10 minutes per side), sonicated for 45 minutes in 3% Hellmanex (Hellma) cleaning solution, washed extensively with ultrapure water, and dried under N_2_ gas. Solutions were placed inside silicone perfusion chambers (CoverWell) pressed on top of the clean glass slides. Prior to each measurement, perfusion chambers were treated with BSA (0.2 mg/mL) for 10 minutes to passivate the glass surface, and then allowed to dry for an additional 10 minutes before filling with the protein solution of interest. Chambers were reused between measurements and rinsed with ultrapure water between samples.

The purity of water used throughout the experiment is critical. All water used for rinsing chambers, preparing background measurements, and making dilutions must be ultrapure (resistivity 18.2 MΩ cm). We have consistently observed an increase in background fluorescence count rate in water stored at room temperature for several days, the origin of which is unclear but may reflect bacterial growth. Water should therefore be used fresh and any unused portion discarded at the end of the day. For the same reason, we store all buffers at −20 °C and thaw them immediately before use, discarding any unused buffer at the end of the experimental session.

### 4.4 Calibration of the Observation Volume

The confocal observation volume is characterized by its radial semiaxis *r*_0_, which appears in the definition of the diffusion time 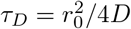 and therefore affects all measured diffusion times. As described in Section 3.4, the fitting procedure is based on the normalized ratio *τ*_app_*/τ*_3_, in which *r*_0_ cancels exactly, so an absolute determination of *r*_0_ is not required for our analysis. The calibration measurement serves instead to monitor the stability of the observation volume throughout the experiment, ensuring that *τ*_app_ values measured at different times can be compared reliably.

Calibration was performed by measuring the autocorrelation decay of a 10 nM solution of TMR in ultrapure water. TMR was chosen as a calibration standard because it is photostable, its absorption maximum is well matched to the 532-nm excitation wavelength, and under our experimental conditions its autocorrelation decay is well described by the single-species diffusion model of Equation 4 with no detectable contributions from triplet blinking or other photophysical processes at short lag times [29]. The autocorrelation decay was fitted with Equation to obtain *τ*_TMR_. In our instrument, *τ*_TMR_ ≈ 100 µs, from which *r*_0_ is calculated as 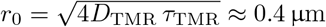 using *D*_TMR_ = 420 µm^2^s^−1^ [39, 40].

### 4.5 Determination of *τ*_*m*_

The diffusion time of the intact oligomer, *τ*_*m*_, is required for the normalization of the titration curves as described in Section 3.4. For PCNA, *τ*_*m*_ = *τ*_3_ cannot be determined from equilibrated samples because measurable dissociation persists even at the highest protein concentration used in our experiments (1 µM PCNA), as shown in Figure 3. Instead, *τ*_3_ is determined by exploiting the slow dissociation kinetics of the trimer: a concentrated stock is diluted to a low PCNA concentration and *τ*_app_ is measured immediately, before significant dissociation has had time to occur. Under these conditions *τ*_app_ ≈ *τ*_3_ regardless of the protein concentration, as illustrated in Figure 3. The mean value of *τ*_app_ obtained from measurements at several concentrations is used as *τ*_3_.

### 4.6 Sample Preparation for Titration Experiments

The goal of the titration experiment is to measure *τ*_app_ over a concentration range that spans the transition between the fully assembled and fully dissociated states of the protein, ideally covering concentrations both well above and well below *K*_*d*_. Because this transition spans several orders of magnitude in concentration, protein concentrations should be distributed approximately logarithmically across the range. If the order of magnitude of the value of *K*_*d*_ is not known in advance, a preliminary experiment covering a wide concentration range can be used to identify the region where *τ*_app_ changes most steeply, and subsequent experiments can sample this region more densely.

A useful quantitative guide for designing the concentration range is provided by the concept of the *logarithmic span* introduced by G. Weber [41]. The logarithmic span is defined as the number of decades in protein concentration over which *α* changes from 0.1 to 0.9, corresponding to the transition from 90% assembled to 90% dissociated. For a homodimer, the equilibrium expression can be rearranged to give *C*_0_*/K*_*d*_ = (1 − *α*)*/*(2*α*^2^). For *α* = 0.1, corresponding to 90% assembled protein, *C*_0_*/K*_*d*_ = 0.9*/*[2(0.1)^2^] = 45, whereas for *α* = 0.9, corresponding to 90% dissociated protein, *C*_0_*/K*_*d*_ = 0.1*/*[2(0.9)^2^] = 1*/*16.2. The logarithmic span is therefore log_10_[45*/*(1*/*16.2)] = log_10_(729) ≈ 2.86 decades in *C*_0_*/K*_*d*_. Thus, to observe the full transition from fully assembled to fully dissociated, a titration for a dimeric protein should span at least three decades of concentration centered around *K*_*d*_.

For a homotrimer (Equation 16), the equilibrium expression can be rearranged to give 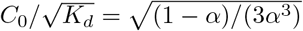. Using the same criterion as above, *α* = 0.1 corresponds to 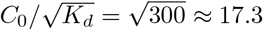, whereas *α* = 0.9 corresponds to 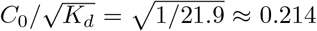. The logarithmic span is therefore log_10_(17.3*/*0.214) = log_10_(81) ≈ 1.9 decades. The logarithmic span also provides a useful diagnostic criterion. An experimental titration curve that appears steeper than predicted by the corresponding equilibrium expression, meaning that it spans fewer decades than expected, may indicate that the assumed dissociation mechanism is incorrect or that an additional process contributes to the observed behavior.

The concentration of labeled protein *C*_*L*_ is kept fixed throughout the titration and should be low enough that the number of labeled particles in the observation volume remains in the optimal range for FCS detection. Concentrations of labeled protein between 1 and 10 nM are typical; we use 1 nM labeled PCNA (*C*_*L*_ = 3 nM) throughout. The total protein concentration *C*_0_ is set by adding the appropriate amount of unlabeled wild-type protein to each sample, so that *C*_0_ = *C*_*L*_ + *C*_*U*_ where *C*_*U*_ is the concentration of unlabeled protein.

An important practical consideration is that the labeled and unlabeled subunits must be allowed to exchange and reach equilibrium before measurements are made. For proteins with slow subunit exchange, this equilibration step can be the most time-consuming part of the experiment. The equilibration time can be assessed by diluting the protein to a concentration where significant dissociation is expected and monitoring *τ*_app_ as a function of time until it reaches a stable plateau. For stable oligomers, equilibration times of tens of hours or more may be required. Insufficient equilibration leads to *τ*_app_ values that do not reflect the true equilibrium distribution of species and will cause *K*_*d*_ to be underestimated.

For the PCNA experiments described here, all titration experiments were performed in a buffer consisting of 20 mM TRIS-HCl (pH 7.3) and 0.1 mg/mL BSA, with a variable concentration of KCl as indicated. Each sample contained 1 nM labeled PCNA-TMR (*f* = 0.6 ± 0.1) and a variable concentration of unlabeled wild-type PCNA to achieve the desired total PCNA concentration, ranging from 1 nM to 1.0 µM. Stock PCNA-TMR was first diluted to a 1 µM PCNA intermediate in 50 mM NaCl, 20 mM TRIS-HCl (pH 7.3), and 0.1 mg/mL BSA, and then further diluted into the sample buffer to a final concentration of 1 nM labeled PCNA. For total PCNA concentrations between 2 and 20 nM, unlabeled wild-type PCNA was first diluted to a 1 µM PCNA intermediate in the same buffer as the labeled intermediate before further dilution into the sample. For total PCNA concentrations greater than 20 nM, the unlabeled stock was diluted directly into the sample. A minimum volume of 500 µL was prepared for each total protein concentration and aliquoted into three semi-independent trials.

Samples were incubated at room temperature for at least 48 hours to ensure equilibration, a time established by monitoring *τ*_app_ as a function of incubation time at a concentration where significant dissociation was expected. Approximately 50 µL of each sample was loaded into the BSA-passivated perfusion chamber for measurement.

Control experiments confirmed that addition of up to 1.0 µM unlabeled wild-type PCNA to a 10 nM TMR solution produced less than 0.5% change in *τ*_TMR_, confirming that protein concentrations in this range do not produce a measurable change in solution viscosity and that changes in *τ*_app_ can be attributed solely to changes in oligomeric state.

A single titration data set provides a preliminary estimate of *K*_*d*_ but is generally insufficient to establish a reliable uncertainty. We recommend collecting at least three independent titration data sets, where each set is prepared and measured on a different day and treated as a fully independent experiment. The *K*_*d*_ values obtained from the individual sets can then be averaged and the standard deviation used as a measure of reproducibility. The number of independent titration sets required depends on the goals of the study. If the aim is simply to obtain a reliable estimate of *K*_*d*_ under a single buffer condition, three to four independent sets are generally sufficient. In our work, where we aim to compare *K*_*d*_ values across multiple co-solute conditions to assess whether they stabilize or destabilize the oligomer, the uncertainty in each *K*_*d*_ must be small enough to support such comparisons and we typically collect more. It is also not uncommon for an entire day of data to be inconsistent with the rest for reasons that are not always identifiable; having additional sets provides the redundancy needed to identify and discard such outliers without compromising the final result.

### 4.7 Data Collection

FCS measurements were carried out using the home-built confocal setup described in Section 4.1, with the 532-nm laser attenuated to a power of 100–120 µW before the entrance of the objective. Before beginning data collection, the instrument is checked and realigned if necessary by loading a 10 nM TMR solution in water into the chamber, acquiring an FCS decay, and fitting to Equation 4 to evaluate the count rate and *r*_0_.

Once the instrument has been checked, background measurements are performed in the same chamber that will be used for the protein samples. A measurement with pure water is used to assess the chamber background and provides the reference autocorrelation function needed for afterpulse correction as described in Section 4.8. The count rate of the water sample establishes a baseline for the instrument background under the specific optical and chamber conditions used; each laboratory should determine their own acceptable threshold by measuring ultrapure water in a clean chamber under their typical operating conditions. In our instrument, count rates typically do not exceed 700 Hz, and the autocorrelation decay does not have features characteristic of diffusing fluorescent particles. Higher count rates usually indicate residual fluorescent contamination requiring additional rinsing of the chamber. For buffer samples, the count rate can be expected to be somewhat higher than that of pure water (up to 2 kHz in our instrument). However, the autocorrelation function should show no evidence of fluorescent contaminants, such as a correlation amplitude above the baseline at short lag times followed by a decay on the characteristic diffusion timescale.

Protein samples are loaded into the chamber sequentially and an FCS decay is acquired as ten consecutive 30-second traces, for a total acquisition time of 5 minutes per sample. Acquiring multiple short traces rather than a single long acquisition is an important practical choice. Protein samples occasionally produce transient spikes in fluorescence intensity, likely due to the passage of large aggregates through the observation volume, which severely distort the autocorrelation function. A single 5-minute acquisition would be entirely compromised by such an event, whereas with 30-second traces only the affected trace needs to be discarded and the remaining traces averaged normally. The frequency with which traces must be discarded varies considerably between protein systems: PCNA behaved very well under the conditions studied here and discarding individual traces was rare, but we have worked with other proteins for which discarding one or two traces per set of 10 was common. The criteria used to identify and remove distorted traces are described in Section 4.9.

To monitor the stability of the laser alignment throughout the experiment, a 10 nM TMR in water sample is measured every 4–6 hours during data collection, and after the final measurement. In our experience, *τ*_TMR_ values measured at different times within the same session agree to within 2%, though differences up to 5% are acceptable. Larger drifts, or any mid-experiment realignment or coverslip replacement, require all background measurements to be repeated before data collection can continue. Data acquired before and after such an interruption are collected under different optical conditions and cannot be normalized using the same *τ*_3_ value; we typically use *τ*_TMR_ measured on both sides of the interruption to correct for the change in *r*_0_ and bring the two sets onto a common scale.

### 4.8 Afterpulse Correction

Avalanche photodiode detectors, which are commonly used in FCS instruments, are subject to an artifact known as afterpulsing. An afterpulse occurs when the detection of a real photon triggers a secondary spurious pulse at the detector shortly afterward, in the absence of a second photon. Because the afterpulse is temporally correlated with the original detection event, it produces a spurious peak in the autocorrelation function at short lag times, distorting the shape of the decay and potentially affecting the measured value of *τ*_app_.

One approach to eliminate afterpulsing is to split the fluorescence signal after the pinhole using a beam splitter and compute the cross-correlation between two independent detectors rather than the autocorrelation of a single channel. Because afterpulse events are not correlated between detectors, the cross-correlation curve is free of this artifact [42]. However, splitting the signal between two detectors reduces the count rate on each detector by approximately half, which decreases the signal-to-noise ratio of the measurement. For single-detector instruments, or when maximizing signal-to-noise is a priority, afterpulsing can instead be removed by a mathematical correction applied to the measured autocorrelation function. Zhao et al. demonstrated that under typical FCS conditions the afterpulsing pattern can be determined from the autocorrelation function of pure water, which contains no fluorescent molecules and therefore provides a non-correlated reference [42]. The corrected autocorrelation function is then obtained as

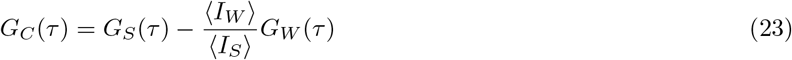

where *G*_*S*_(*τ*) is the measured autocorrelation function of the sample, *G*_*W*_ (*τ*) is the autocorrelation function of pure water measured under identical conditions, *G*_*C*_(*τ*) is the corrected autocorrelation function, and ⟨*I*_*W*_ ⟩ and ⟨*I*_*S*_⟩ are the mean count rates of the water and sample measurements, respectively. The scaling factor ⟨*I*_*W*_⟩ */* ⟨*I*_*S*_⟩ accounts for the difference in count rate between the water reference and the protein sample, as required by the derivation of Zhao et al. [42]. This correction is valid provided that the afterpulsing probability is small and its characteristic decay time is much shorter than the diffusion time of the fluorescent particles, conditions that are well satisfied in typical FCS instruments. An example of the correction applied to an experimental PCNA autocorrelation decay is shown in Figure 5, where the spurious contribution at short lag times is clearly visible in the uncorrected decay and is completely removed after applying Equation 23. The correction is applied to each individual 30-second autocorrelation trace before the traces are averaged.

**Figure 5:**
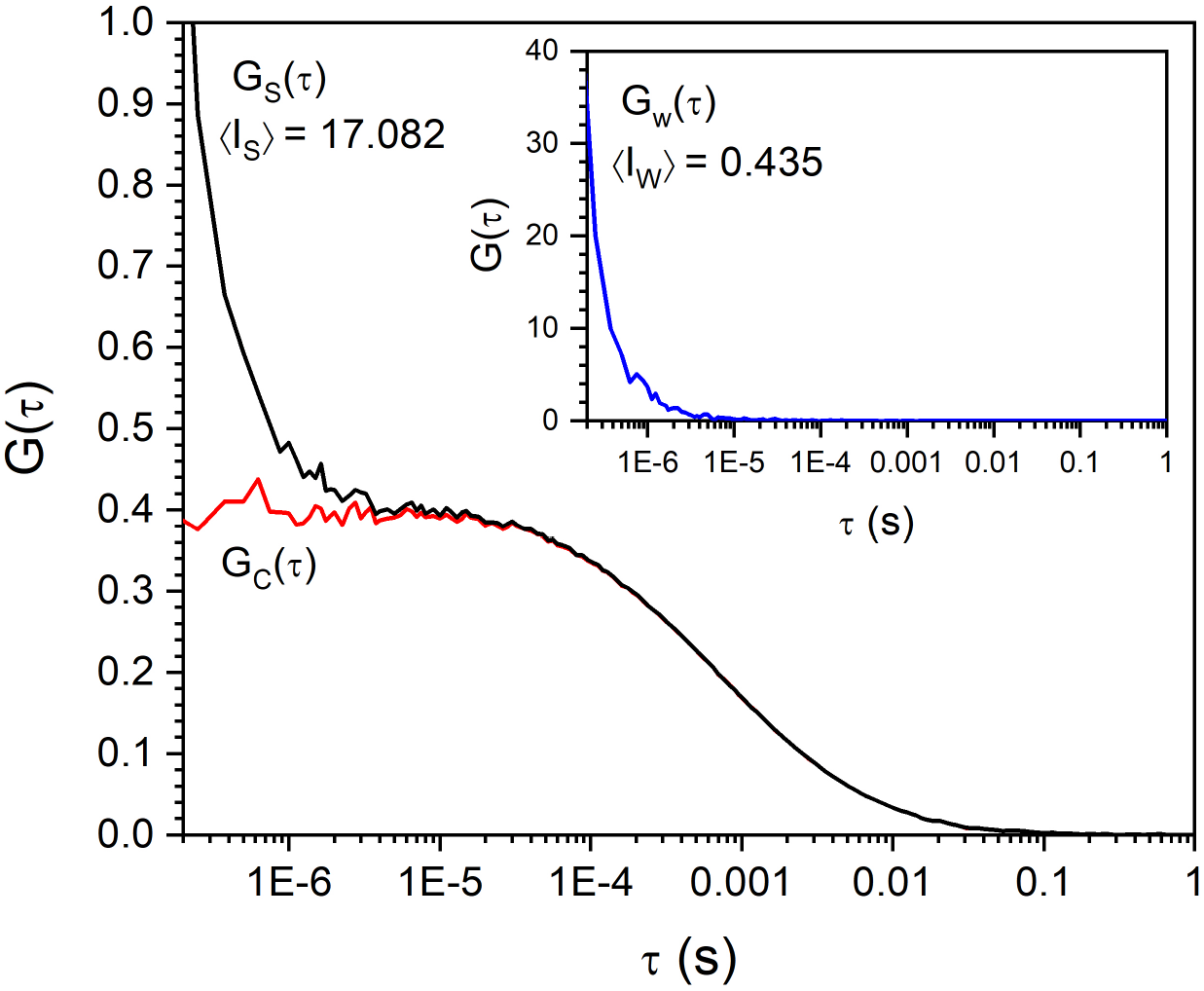
Illustration of the afterpulse correction procedure. The main panel shows the measured autocorrelation function of a PCNA sample, *G*_*S*_(*τ*) (black), and the corrected autocorrelation function *G*_*C*_ (*τ*) (red), obtained using Equation 23. The afterpulse artifact is visible as a large spurious contribution at short lag times (*τ <* 10^−5^ s) in *G*_*S*_(*τ*), which is absent in *G*_*C*_ (*τ*). The two curves overlap at longer lag times where the afterpulse contribution is negligible. The inset shows the autocorrelation function of pure water, *G*_*W*_ (*τ*), which serves as the non-correlated reference for the correction. The large amplitude and rapid decay of *G*_*W*_ (*τ*) reflect the afterpulsing characteristics of the detector; the mean count rates ⟨*I*_*S*_⟩ = 17.08 kHz and ⟨*I*_*W*_⟩ = 0.44 kHz are used to scale the water reference before subtraction as described in Equation 23.

### 4.9 Quality Control and Outlier Removal

After afterpulse correction, the individual 30-second traces are evaluated before averaging. Each corrected trace is fitted individually with Equation 4 to obtain a *τ*_app_ value, and the median *τ*_app_ across all ten traces is computed. Any trace whose *τ*_app_ deviates from the median by more than 15% is excluded from the average. In our experience, this threshold is sufficient to remove clearly distorted traces, typically caused by transient spikes in fluorescence intensity from aggregates or dust particles passing through the observation volume, without discarding traces that are otherwise acceptable. Figure 6 (left) shows a representative example in which one of ten traces is identified and removed by this criterion. The practical value of this approach is that it allows the entire analysis pipeline – afterpulse correction, outlier removal, and averaging – to be automated, producing a final average autocorrelation function ready for fitting without requiring case-by-case visual inspection of individual traces.

**Figure 6:**
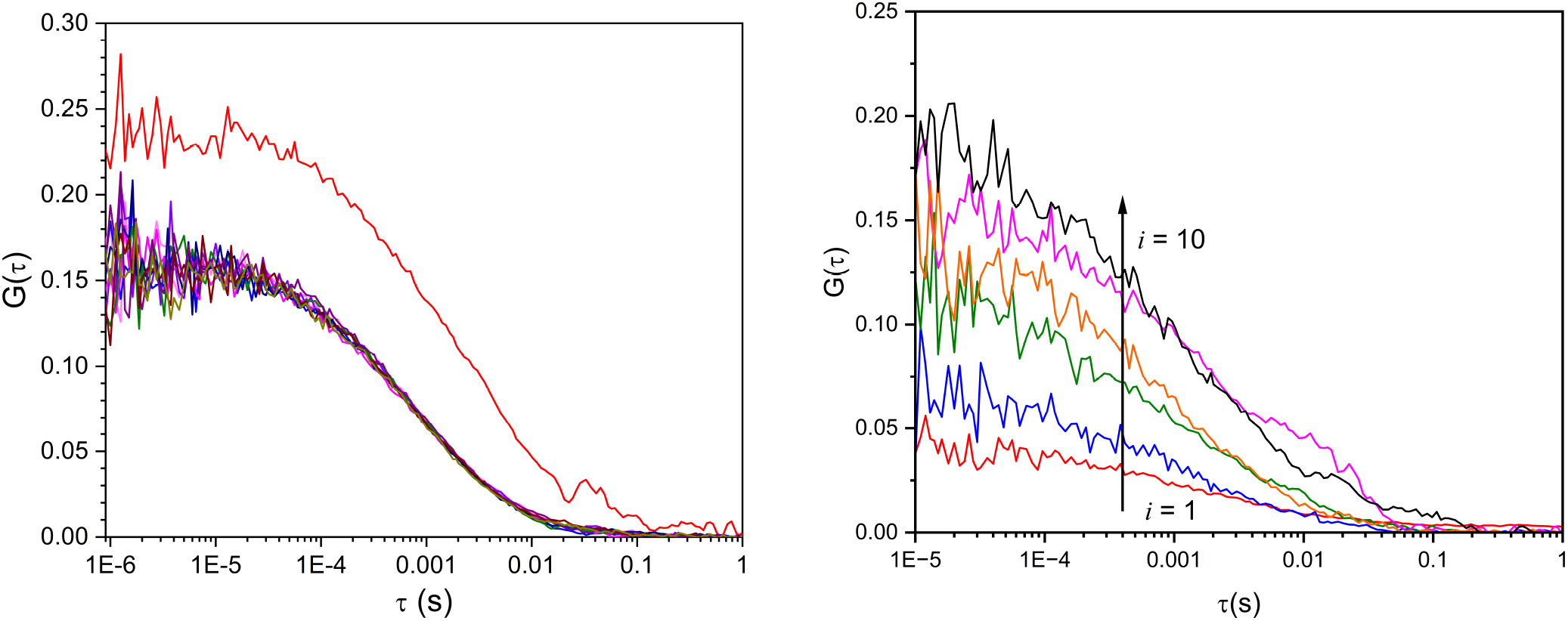
Left: Representative set of ten individual 30-second autocorrelation traces acquired for a single PCNA sample, illustrating the outlier removal procedure described in Section 4.9. One trace (red) is identified as an outlier based on its *τ*_app_ deviating by more than 15% from the median, most likely due to a transient spike in fluorescence intensity from an aggregate passing through the observation volume. This trace is excluded and the remaining nine traces are averaged to produce the final autocorrelation function used for fitting. The distortion in the outlier trace is most visible at longer lag times, which is a characteristic signature of a fluorescence intensity spike during acquisition. Right: Example of a systematic drift in *G*_0_ across ten consecutive 30-second traces. The sample contained an aggregation-prone protein for which BSA passivation was insufficient. The amplitude *G*_0_ increases approximately fourfold from the first to the last trace while the fluorescence intensity decreases by approximately the same factor (not shown). This behavior is consistent with progressive loss of fluorescent particles from solution due to surface adsorption (see Section 4.9 for discussion).

In addition, the amplitude *G*_0_ and the count rate are monitored throughout data collection as real-time indicators of sample and instrument stability. A systematic drift in *G*_0_ from the first to the last trace within a single sample is a clear warning sign. Figure 6 (right) shows an example in which *G*_0_ increases approximately fourfold over ten consecutive 30-second traces, indicating a progressive loss of labeled protein from solution most likely due to adsorption onto the chamber surface. This behavior signals that the surface passivation is insufficient for the protein under study and the protocol should be revised before proceeding with quantitative measurements.

We emphasize that none of the criteria described above are strict rules. The thresholds and procedures described here reflect the conventions adopted in our laboratory for PCNA and related systems, and are intended as a practical starting point rather than a prescriptive protocol. Researchers should exercise their own judgment based on their knowledge of the protein system and the overall quality of the data, and whatever criteria are adopted should be applied consistently across all samples within a dataset.

## 5 Results

### 5.1 Titration Curves and Determination of *K*_*d*_

To illustrate the application of the method described in the preceding sections, we present experimental data for *S. cerevisiae* PCNA in buffers containing KCl at concentrations ranging from 50 mM to 1.0 M, prepared in 20 mM TRIS-HCl (pH 7.3) and 0.1 mg/mL BSA. These experiments were designed to assess the effect of ionic strength on the stability of the PCNA trimer. The entire experiment, including thawing the stock solution, preparing the dilution series, incubating the samples, and acquiring the FCS data, was repeated independently on three different days. Within each independent experiment, three separate aliquots were prepared from each final dilution, and each aliquot was measured once by FCS. Therefore, each concentration point was characterized by three technical replicates in each of three independent experiments, yielding a total of nine measurements per concentration.

Figure 7 shows a representative titration curve for PCNA in 250 mM KCl. The normalized apparent diffusion time *τ*_app_*/τ*_3_ decreases monotonically as the total protein concentration *C*_0_ is reduced, reflecting the progressive dissociation of trimers into monomers at lower concentrations. At high protein concentrations, *τ*_app_*/τ*_3_ approaches unity, consistent with a solution dominated by intact trimers. At low concentrations, *τ*_app_*/τ*_3_ approaches 3^−1*/*3^ ≈ 0.69, the value expected for a solution of free monomers. Each data point represents the average of all independent titration sets at that concentration, and the error bars are 95% confidence intervals computed using a *t*-distribution multiplier appropriate for the number of measurements available. The solid line shows the fit of Equations 16 and 21 to the data, with *K*_*d*_ as the only free parameter and *τ*_1_*/τ*_3_ fixed at 3^−1*/*3^, weighted by the inverse of the 95% confidence intervals. Using confidence intervals rather than standard deviations as weights places greater emphasis on concentration points with more replicates, which is appropriate when the number of independent sets varies across the titration curve; researchers may choose to use standard deviations instead with little effect on the extracted *K*_*d*_. The fit describes the data well across the entire concentration range, validating the underlying physical model.

**Figure 7:**
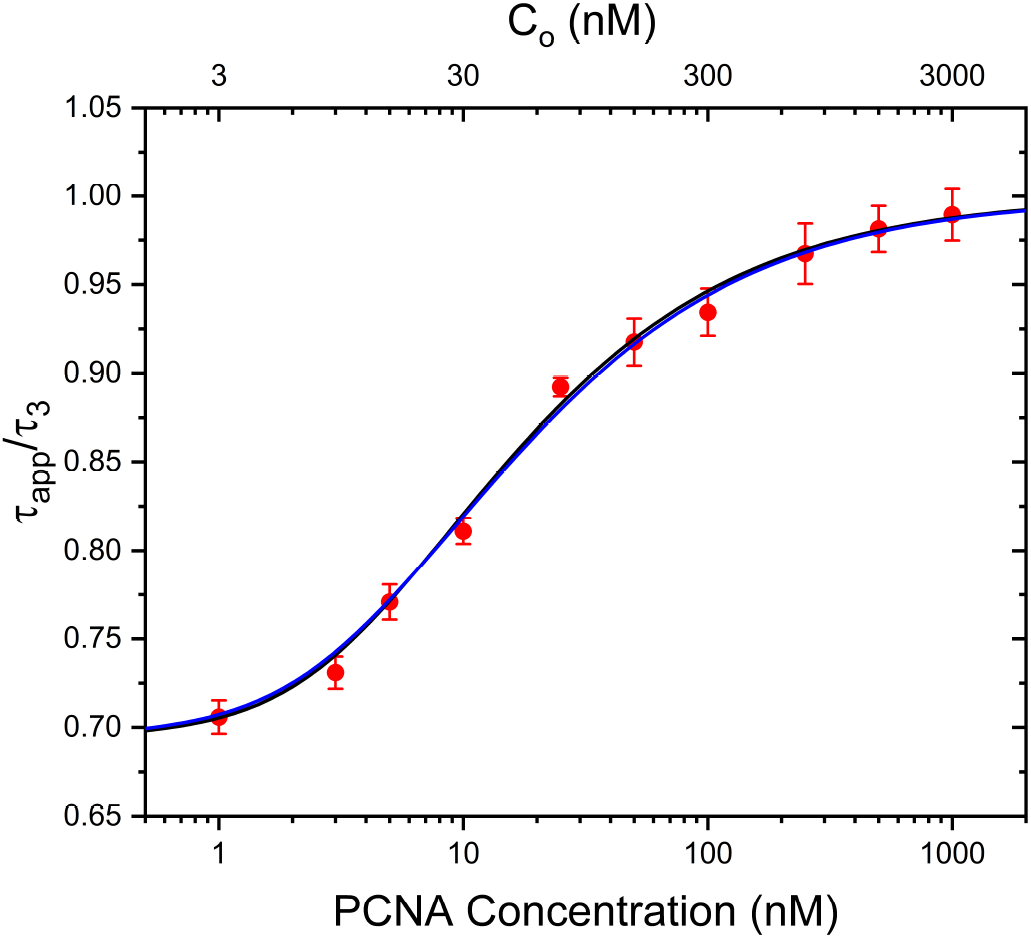
Representative FCS titration curve for PCNA in 250 mM KCl, 20 mM TRIS-HCl (pH 7.3), and 0.1 mg/mL BSA. The normalized apparent diffusion time *τ*_app_*/τ*_3_ is plotted as a function of total PCNA concentration 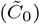. The top axis shows the corresponding total protein concentration expressed in terms of monomers (*C*_0_). Red circles are experimental data points with error bars representing 95% confidence intervals. The black and blue curves show the best fits of Equations S2 and 21 to the data with labeling efficiencies *f* = 0.6 and *f* = 1.0, respectively. The two fits are nearly indistinguishable, demonstrating the insensitivity of the extracted *K*_*d*_ to the precise value of *f*.

Figure 8 shows titration curves for two representative KCl concentrations, 50 mM and 500 mM, illustrating the effect of salt on PCNA trimer stability. The curve shifts to lower PCNA concentrations as KCl increases from 50 to 500 mM, indicating that the trimer is more stable at higher salt concentration.

**Figure 8:**
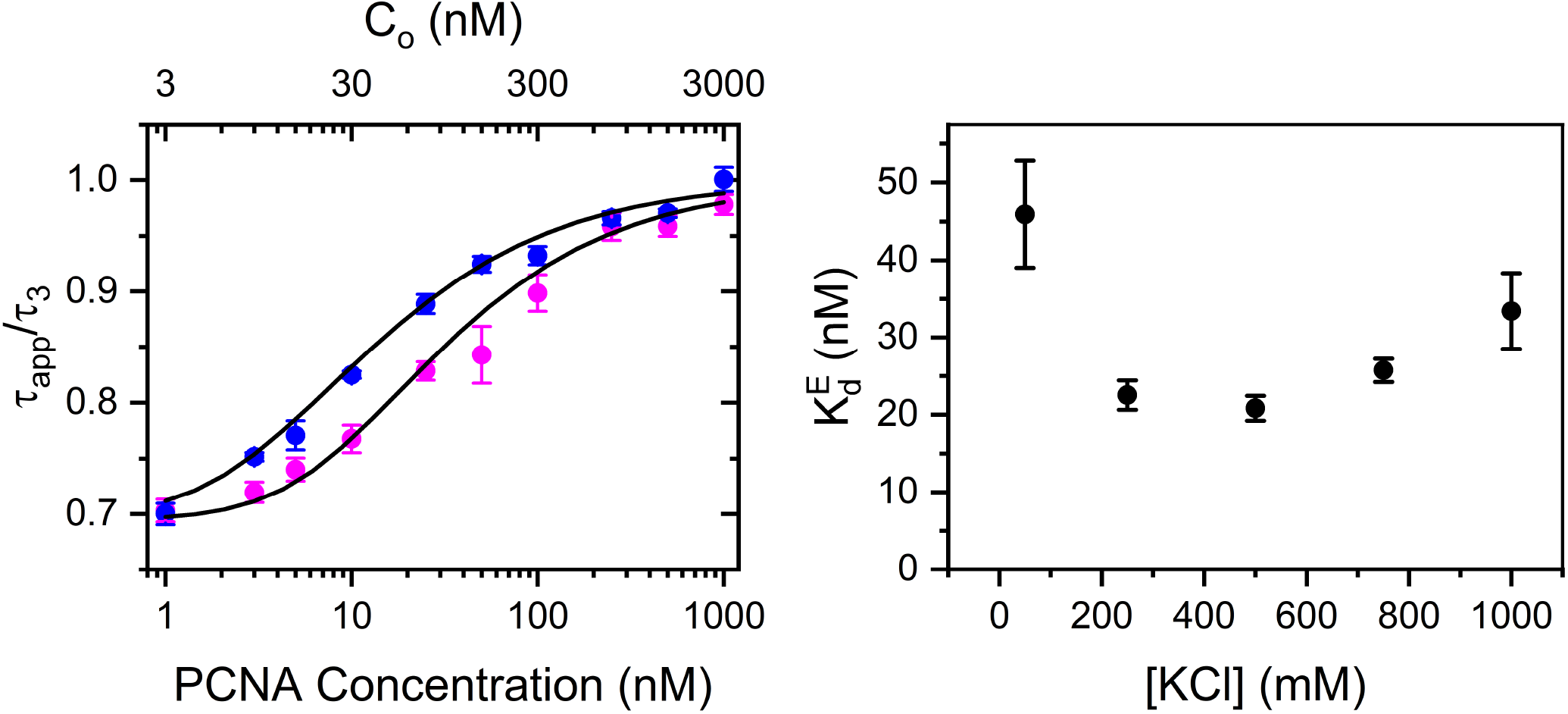
Effect of [KCl] on PCNA trimer stability. (Left) Representative FCS titration curves for PCNA in 50 mM KCl (blue) and 500 mM KCl (magenta), both in 20 mM TRIS-HCl (pH 7.3) and 0.1 mg/mL BSA. The normalized apparent diffusion time *τ*_app_*/τ*_3_ is plotted as a function of total PCNA concentration 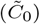. The top axis shows the corresponding total protein concentration expressed in terms of monomers (*C*_0_). Solid curves show the best fits of Equations S2 and 21 to the data. (Right) Effective dissociation concentration 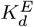 as a function of KCl concentration, calculated from the fitted *K*_*d*_ values using Equation 24. Error bars represent 95% confidence intervals. 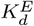 reaches a minimum in the 250–500 mM range, indicating that moderate KCl stabilizes the PCNA trimer, while very high KCl concentrations partially destabilize the complex.

Because *K*_*d*_ for a trimer-monomer equilibrium has units of concentration squared (nM^2^), it is not directly interpretable as a concentration. To facilitate comparison with dissociation constants from other oligomerization mechanisms, we also report an effective dissociation concentration 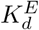, defined as the total protein concentration *C*_0_, expressed in terms of monomers, at which half of all protein subunits are present as free monomers and half are incorporated into trimers (*α* = 1*/*2). Setting *α* = 1*/*2 in Equation 15 and solving for *C*_0_ gives (see Supplemental Information file)

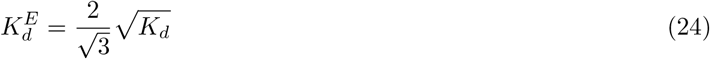

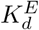 is expressed in units of nM and is analogous to the *K*_*d*_ of a standard 1:1 binding equilibrium in the sense that it defines the concentration scale over which the transition between associated and dissociated states occurs. Both *K*_*d*_ and 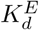 are reported in Table 1.

**Table 1:** Equilibrium dissociation constants for PCNA in KCl buffers containing 20 mM TRIS-HCl (pH 7.3) and 0.1 mg/mL BSA. *K*_*d*_ values were obtained from nonlinear least-squares fits of the normalized titration curves. The effective dissociation concentration 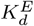 was calculated from *K*_*d*_ using Equation 24.

| [KCl] (mM) | $K_d$ (nM<sup>2</sup>) | $K_d^E$ (nM) |
| --- | --- | --- |
| 50 | $(4.7 \pm 1.4) \times 10^3$ | $79 \pm 12$ |
| 250 | $(1.14 \pm 0.18) \times 10^3$ | $39.0 \pm 3.1$ |
| 500 | $(0.99 \pm 0.18) \times 10^3$ | $36.3 \pm 3.3$ |
| 750 | $(1.48 \pm 0.17) \times 10^3$ | $44.4 \pm 2.6$ |
| 1000 | $(2.50 \pm 0.72) \times 10^3$ | $57.7 \pm 8.3$ |

The *K*_*d*_ values obtained from fitting the titration curves measured at five KCl concentrations are listed in Table 1 and the corresponding 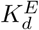 values are plotted as a function of KCl concentration in Figure 8 (right panel). The full dataset reveals that stabilization is most pronounced between 50 and 250 mM KCl, and that the trend reverses at higher concentrations, with 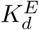 increasing again above 500 mM KCl, suggesting that very high salt concentrations partially destabilize the oligomeric complex.

The data reveal a clear dependence of PCNA trimer stability on KCl concentration. The 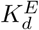 value drops from approximately 79 nM at 50 mM KCl to a minimum of approximately 36–39 nM in the 250–500 mM range, indicating that moderate salt concentrations stabilize the trimer. At higher KCl concentrations 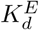 increases again, reaching approximately 58 nM at 1.0 M KCl, suggesting that very high salt concentration partially destabilizes the oligomeric complex. The observed stabilization of the trimer at intermediate salt concentrations likely reflects electrostatic screening of repulsive interactions at the subunit interfaces, whereas at higher salt concentrations this effect is offset by nonspecific ionic interactions that reduce trimer stability.

### 5.2 Insensitivity of *K*_*d*_ to Labeling Efficiency

As noted in Section 3.4, the extracted *K*_*d*_ depends in principle on the labeling efficiency *f* through the parameter *p* = *f C*_*L*_*/C*_0_. In practice, however, *K*_*d*_ is relatively insensitive to the precise value of *f*, which has important practical implications: an approximate determination of *f* is sufficient for reliable results, and systematic errors in the measurement of *f* do not strongly affect the conclusions drawn from the titration data.

This insensitivity is illustrated in Figure 7, which shows fits to the 250 mM KCl titration data obtained with *f* = 0.6 (black) and *f* = 1.0 (blue). Fitting the same experimental data with *f* = 0.6 and *f* = 1.0 yields *K*_*d*_ values of (11.4 ± 1.8) × 10^2^ nM^2^ and (13.4 ± 2.1) × 10^2^ nM^2^, respectively, which differ by less than 20% and are well within their mutual uncertainties. The fitted curves for the two labeling efficiencies are visually indistinguishable. This insensitivity arises because *p* = *f C*_*L*_*/C*_0_ is small throughout most of the titration. Since the labeled PCNA concentration 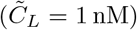 is much smaller than *C*_0_ at most concentration points, the precise value of *f* has little influence on the predicted *τ*_app_.

## 6 Extensions and Limitations

The method described in this work is broadly applicable to homomeric protein complexes, but several practical considerations define the range of systems and conditions for which it is suitable.

### 6.1 Accessible Range of *K*_*d*_

The approach described here for the determination of *τ*_3_ relies on the slow dissociation kinetics of the intact oligomer: a freshly diluted protein must remain essentially fully assembled on the timescale of the measurement. This condition is satisfied when the dissociation rate constant is small, which is generally the case for stable oligomers with small *K*_*d*_ values. For less stable complexes, dissociation may be fast enough that *τ*_app_ begins to decrease before the first measurement can be acquired, making it impossible to determine *τ*_*m*_ by this approach. In such cases, *τ*_*m*_ must be determined at sufficiently high protein concentration that dissociation is negligible at equilibrium. FCS measurements at high protein concentrations require working with a small concentration of labeled protein (section 2.3) to keep the number of fluorescent particles in the observation volume within the optimal range for FCS detection. In our experience, total protein concentrations up to approximately 100 µM are accessible under these conditions, as demonstrated in our studies of RuBisCO activase [14], though the practical upper limit will depend on the solubility and aggregation properties of the specific protein under study. The accessible concentration range sets an upper bound on the 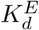 values that can be determined, which in practice is at least one order of magnitude below the highest accessible concentration.

At the other extreme, if the complex is extremely stable and 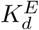 falls well below the lowest accessible protein concentration, dissociation occurs outside the measurable range and the titration curve will show no measurable change in *τ*_app_. In this case, only a lower bound on 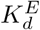 can be established. The lowest concentration at which reliable FCS measurements can be acquired is determined by the signal-to-noise ratio of the autocorrelation decay. In our instrument, measurements are routinely performed at 1 nM labeled protein and can be pushed to approximately 100 pM, below which the signal-to-noise ratio becomes too poor for reliable fitting.

### 6.2 Two-State Dissociation

The analysis presented here assumes that the protein undergoes a two-state transition between the intact oligomer and free monomers, with no accumulation of partially dissociated intermediates. For PCNA, this assumption is supported by experimental evidence [13]. For other trimeric or higher-order oligomers, the presence of intermediates should be assessed independently, for example by monitoring whether the shape of the titration curve is consistent with the two-state model. A systematic deviation of the experimental data from the fitted curve may indicate the presence of intermediates or a more complex dissociation mechanism.

### 6.3 Broader Applications and Simulation Tool

The mathematical framework presented here for a homotrimeric protein is readily adapted to other oligomerization stoichiometries. The dimer case is analytically simpler, with *α* given by the closed-form expression in Equation S1. The tetramer case, including the more complex situation where dissociation proceeds via a dimeric intermediate, is discussed in detail in Kanno and Levitus [19]. To assist researchers in designing titration experiments and evaluating the expected behavior of their protein of interest, we have developed an online simulation tool using Streamlit (https://protein-oligomerization-equilibria-fna67mqcnzwacp7iok5uvb.streamlit.app). The app can also run locally using the instructions detailed in GitHub (https://github.com/ddrathod-create/FCS-Protein-Oligomerization-Equilibria). The shape and position of the titration curve depend on *K*_*d*_, *C*_*L*_, and *f* in a non-trivial way, and it is not always intuitive whether a given *K*_*d*_ will produce a measurable change in *τ*_app_ over an experimentally accessible concentration range. The tool allows users to simulate titration curves for three oligomerization mechanisms – dimer-monomer, trimer-monomer, and sequential tetramer-dimer-monomer – by inputting *K*_*d*_ values and experimental parameters (*C*_*L*_, *f*). The tool computes and displays the predicted normalized apparent diffusion time *τ*_*n*_ = *τ*_app_*/τ*_*m*_ as a function of total protein concentration, allowing researchers to assess whether a given *K*_*d*_ is accessible within their experimental concentration range and what concentration range is needed to adequately sample the transition between assembled and dissociated states before committing to a full set of experiments. An example of simulated titration curve generated with the tool is provided in the Supplemental Information file.

The same mathematical framework also extends naturally to the determination of dissociation and association rate constants from time-dependent *τ*_app_ measurements, in which the protein is diluted from a concentrated stock and FCS decays are acquired as a function of time as the system approaches equilibrium. This approach, and the analytical expressions that relate the time-dependent *τ*_app_ values to the kinetic rate constants, are described in Kanno and Levitus [19] and have been applied in our laboratory to characterize the dissociation and association kinetics of PCNA and the *E. coli β*-clamp [9, 13]. A detailed treatment of kinetic experiments is beyond the scope of this article.

## 7 Conclusions

We have presented a comprehensive practical guide to the determination of protein oligomerization equilibrium constants by fluorescence correlation spectroscopy. Using the homotrimeric sliding clamp PCNA as a model system, we have described the complete experimental workflow from instrument calibration and sample preparation through data collection, quality control, and fitting, illustrating each step with experimental data. The mathematical framework underlying the analysis, originally derived by Kanno and Levitus [19] and generalized here to arbitrary oligomeric stoichiometry, provides an analytically tractable relationship between the experimentally measured apparent diffusion time *τ*_app_ and the dissociation equilibrium constant *K*_*d*_, accounting rigorously for the distribution of fluorescent labels among oligomeric species and the brightness-squared weighting of the autocorrelation function.

Several practical aspects of the method deserve emphasis. The normalization of *τ*_app_ by the diffusion time of the intact oligomer *τ*_*m*_, determined from freshly diluted samples before dissociation has occurred, eliminates the dependence of the results on instrumental parameters and allows direct comparison across experiments performed on different days or in different laboratories. The extracted *K*_*d*_ is relatively insensitive to the labeling efficiency *f*, which means that an approximate determination of *f* is sufficient for reliable results. The method is applicable over a wide range of *K*_*d*_ values, from values somewhat above the lowest accessible protein concentration (approximately 100 pM in our instrument) up to values set by the highest accessible protein concentration, which in our experience extends to tens of micromolar.

The application of the method to PCNA in buffers of varying KCl concentration illustrates its ability to detect and quantify changes in oligomer stability as a function of solution conditions. The effective dissociation concentration 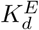 reaches a minimum in the 250–500 mM KCl range, indicating that moderate salt concentrations stabilize the PCNA trimer, while very high KCl concentrations partially destabilize the complex. This type of systematic study, in which *K*_*d*_ is determined under multiple solution conditions and the results compared quantitatively, is one of the most powerful applications of the method.

We anticipate that this practical guide will lower the barrier to implementing quantitative FCS-based oligomerization studies, and will encourage the adoption of physically accurate models in place of empirical approaches that can yield misleading results.

## Acknowledgments

We thank past members of the Levitus lab at Arizona State University who contributed to the development and implementation of FCS instrumentation and analysis methods over the past two decades: Kaushik Gurunathan, Suman Ranjit, Manas Chakraborty, Jennifer England, David Kanno, Andrew Serban, Bryan Donaphon, and Anirban Purohit. We also thank our collaborators Linda Bloom and Rebekka Wachter, who provided the protein systems that enabled the application and validation of these methods.

## Conflict of Interest Statement

The authors have declared that no conflicting interests exist.

## Notes

### Competing Interest Statement

The authors have declared no competing interest.

