## Supplemental Materials for "A practical framework for measuring protein oligomerization equilibria by fluorescence correlation spectroscopy"

### S.1 Definition of $\tau_{\text{app}}$

Experimental autocorrelation decays are fitted with the single-species model of Equation 4,

$$G(\tau) = G_0 \left(1 + \frac{\tau}{\tau_{\text{app}}}\right)^{-1}$$

where $\tau_{\text{app}}$ is the effective diffusion time obtained from fitting the measured decay. At $\tau = \tau_{\text{app}}$, the autocorrelation function has decayed to half its initial value,

$$G(\tau_{\text{app}}) = \frac{G_0}{2}$$

For a mixture containing monomers and a single oligomeric species with stoichiometry $m$, the autocorrelation function is given by Equation 17,

$$G(\tau) = A_1 \left(1 + \frac{\tau}{\tau_1}\right)^{-1} + A_m \left(1 + \frac{\tau}{\tau_m}\right)^{-1}$$

Setting $G(\tau_{\text{app}}) = G_0/2$ and noting that $G_0 = A_1 + A_m$ gives

$$A_1 \left(1 + \frac{\tau_{\text{app}}}{\tau_1}\right)^{-1} + A_m \left(1 + \frac{\tau_{\text{app}}}{\tau_m}\right)^{-1} = \frac{A_1 + A_m}{2}$$

Multiplying through by $2\,(1 + \tau_{\text{app}}/\tau_1)\,(1 + \tau_{\text{app}}/\tau_m)$ and expanding gives

$$2A_1 \left(1 + \frac{\tau_{\text{app}}}{\tau_m}\right) + 2A_m \left(1 + \frac{\tau_{\text{app}}}{\tau_1}\right) = (A_1 + A_m) \left(1 + \frac{\tau_{\text{app}}}{\tau_1}\right) \left(1 + \frac{\tau_{\text{app}}}{\tau_m}\right)$$

Expanding and collecting terms in powers of $\tau_{\text{app}}$ yields a quadratic equation in $\tau_{\text{app}}$,

$$\tau_{\text{app}}^2 - c(\tau_m - \tau_1)\tau_{\text{app}} - \tau_1\tau_m = 0$$

where

$$c = \frac{A_1 - A_m}{A_1 + A_m}$$

### S.2 Generalization to Arbitrary Stoichiometry $m$

It remains to express $c$ in terms of physically meaningful quantities. Using Equation 13, the amplitudes are

$$A_1 = \kappa\,\alpha_1 \qquad A_m = \kappa\,\alpha_m[1 + (m-1)p]$$

where $\alpha_1 = \alpha$ and $\alpha_m = 1 - \alpha$ for a two-species system. Substituting gives

$$c = \frac{\alpha - (1-\alpha)[1 + (m-1)p]}{\alpha + (1-\alpha)[1 + (m-1)p]}$$

Rearranging gives

$$c = 1 - \frac{2\alpha}{1 + (1-\alpha)(m-1)p} \cdot \frac{1 + (1-\alpha)(m-1)p}{\alpha + (1-\alpha)[1 + (m-1)p]}$$

which can be written more compactly as

$$c = 1 - \frac{2\alpha}{1 + (1-\alpha)(m-1)p}$$

which is Equation 19. For $m = 2$ this reduces to $c = 1 - 2\alpha/(1 + (1-\alpha)p)$, which is the expression derived by Kanno and Levitus [1] for the dimer case.

### S.3 Analytical Solution for the Degree of Dissociation of a Homodimeric or Homotrimeric Protein

The positive root of the quadratic equation relating $\alpha$ to $K_d$ and $C_0$ for a homodimeric protein (Equation 14) is

$$\alpha = \frac{1}{4C_0}\left(-K_d + \sqrt{K_d^2 + 8K_dC_0}\right) \tag{S1}$$

The cubic equation relating $\alpha$ to $K_d$ and $C_0$ for a homotrimeric protein (Equation 16) can be solved analytically. Dividing through by $3C_0^2$ and defining the dimensionless parameter $\beta = K_d/3C_0^2$ gives the depressed cubic

$$\alpha^3 + \beta\alpha - \beta = 0$$

whose unique real root in the interval $0 \leq \alpha \leq 1$ is given by Cardano's formula,

$$\alpha = \left[\frac{\beta}{2}\left(1 + \sqrt{1 + \frac{4\beta}{27}}\right)\right]^{1/3} + \left[\frac{\beta}{2}\left(1 - \sqrt{1 + \frac{4\beta}{27}}\right)\right]^{1/3} \tag{S2}$$

The argument of the square root, $1 + 4\beta/27$, is always positive for physical values of $K_d$ and $C_0$, ensuring that the solution is real and unique. The parameter $\beta$ has a useful physical interpretation: when $\beta \ll 1$, corresponding to low $K_d$ or high $C_0$, the protein is predominantly assembled and $\alpha \approx 0$; when $\beta \gg 1$, corresponding to high $K_d$ or low $C_0$, the protein is predominantly dissociated and $\alpha \approx 1$.

### S.4 Derivation of the Effective Dissociation Concentration $K_d^E$

The effective dissociation concentration $K_d^E$ is defined as the total monomer concentration $C_0$ at which half of all protein subunits are present as free monomers, i.e. $\alpha = 1/2$. Substituting $\alpha = 1/2$ into Equation 15 gives

$$K_d = \frac{3(1/2)^3 C_0^2}{1 - 1/2} = \frac{3C_0^2}{8} \cdot 2 = \frac{3C_0^2}{4}$$

Solving for $C_0 \equiv K_d^E$ gives

$$K_d^E = \frac{2}{\sqrt{3}}\sqrt{K_d}$$

which is Equation 24.

### S.5 Simulation Tool

We have developed an online simulation tool using Streamlit:
(https://protein-oligomerization-equilibria-fna67mqcnzwacp7iok5uvb.streamlit.app).
The app can also run locally using the instructions detailed in GitHub (https://github.com/ddrathod-create/FCS-Protein-Oligomerization-Equilibria). This tool was created to allow users to simulate titration curves for three oligomerization mechanisms:

1) dimer ($D$)-monomer ($M$)

$$D \rightleftharpoons 2\,M \quad K_d = \frac{[M]^2}{[D]}$$

2) trimer ($T_3$)-monomer ($M$)

$$T_3 \rightleftharpoons 3\,M \quad K_d = \frac{[M]^3}{[T_3]}, \qquad K_d^E = \frac{2}{\sqrt{3}}\sqrt{K_d}$$

3) tetramer ($T_4$)-dimer ($D$)-monomer ($M$)

$$T_4 \rightleftharpoons 2\,D \quad K_{d1} = \frac{[D]^2}{[T_4]}$$

$$D \rightleftharpoons 2\,M \quad K_{d2} = \frac{[M]^2}{[D]}$$

Here, we use $T_3$ to denote a trimer, and $T_4$ to denote a tetramer. In each case, the inputs are the $K_d$ values ($K_d^E$ for the second case), and experimental parameters: the concentration of labeled protein ($\tilde{C}_L$, expressed in terms of oligomers), and the labeling efficiency ($f$). The tool computes and displays the predicted normalized apparent diffusion time $\tau_n = \tau_{\text{app}}/\tau_m$ as a function of total protein concentration $\tilde{C}_0$.

The example below shows the output for the dissociation of a tetramer with a stable dimeric intermediate.

INPUTS:

$$T_4 \rightleftharpoons 2\,D \quad K_{d1} = \frac{[D]^2}{[T_4]} = 100\,\text{nM}$$

$$D \rightleftharpoons 2\,M \quad K_{d2} = \frac{[M]^2}{[D]} = 10\,\text{nM}$$

Labeling efficiency: $f = 0.5$

Concentration of labeled protein: $\tilde{C}_L = 1\,\text{nM}$ (expressed in terms of tetramers, $C_L = 4\,\text{nM}$).

Total concentration of protein (labeled + unlabeled): $\tilde{C}_0 = 1\,\text{nM} - 1000\,\text{nM}$ (expressed in terms of tetramers, $C_0 = 4\,\text{nM}$).

OUTPUT Top panel: $\tau_{\text{app}}/\tau_4$ vs total protein concentration ($\tilde{C}_0$ in bottom axis, $C_0$ in top axis). $\tau_4$ is the diffusion time of the intact tetramer. Bottom pannel: fractional concentrations ($\alpha_1, \alpha_2, \alpha_4$) vs total protein concentration.

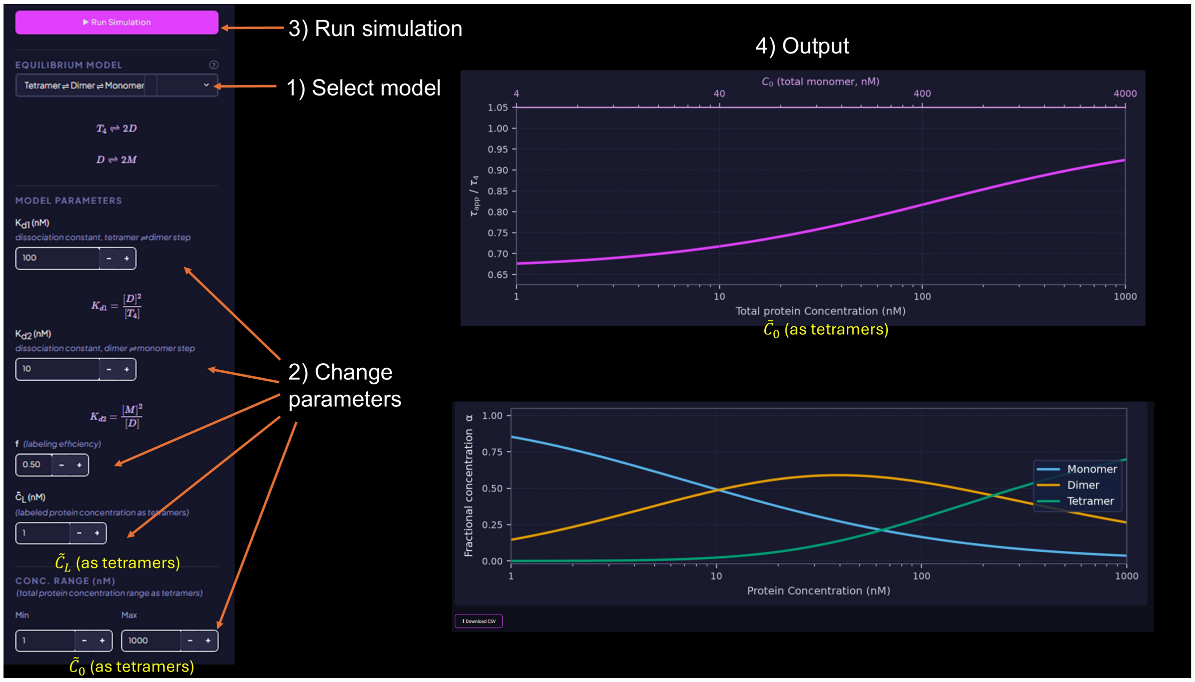


The "download CSV" button generates and downloads a file containing the information of the graphs in CSV format.
